# Attention Improves Population Codes by Warping Neural Manifolds in Human Visual Cortex

**DOI:** 10.1101/2025.10.09.681102

**Authors:** Yu-Qi You, Shurui Li, Yu-Ang Cheng, Yuanning Li, Kendrick Kay, Ru-Yuan Zhang

**Author notes:** Corresponding Ru-Yuan Zhang, Shanghai Jiao Tong University.

## Abstract

Decades of research on visual attention have revealed its numerous effects on neural responses. Two competing models have been proposed for how these effects lead to improved population representations: one highlights changes in neural tuning, while the other points towards changes in trial-by-trial noise correlations. Here, we develop a neural population manifold framework that interprets changes in neural responses as geometric transformations in high-dimensional neural space, allowing us to disentangle and quantify the effects of tuning changes and correlation changes induced by attention. Applying this framework to extensive measurements of cortical responses during different attentional tasks, we find that tuning changes are the primary driver of improved population representations. In contrast, correlation changes, though present, have minimal—or even detrimental—effects to information content due to its strong interactions with other changes (e.g., tuning, variability). Our results support the “tuning change” model of visual attention and demonstrate a general framework for adjudicating how different aspects of neural coding affect information processing.

## INTRODUCTION

Visual inputs are remarkably vast, necessitating efficient selection mechanisms that enable the brain to flexibly process information in response to complex task demands. Over the past few decades, converging evidence from neuroscience, psychology, and artificial intelligence has identified attention as one of the most fundamental selection mechanisms in the brain^1^. While the attentional enhancement of behavioral performance is well-documented in a wide range of visual tasks^2^, our understanding of the underlying neural mechanisms remains limited^3,4^.

Previous human imaging studies have shown that attention enhances the decoding accuracy of stimuli across multiple brain regions, including LGN, V1, V2, and V4^5-7^. Similarly, multi-unit recordings in macaques have demonstrated that attention significantly enhances the encoding of sensory information in population activity within areas such as V4 and MT^8-11^. These converging findings from both humans and macaques provide strong evidence for the view that attention improves behavior by refining population-level neural representations.

But how exactly are those representations improved? Two competing models have been proposed. The “tuning change” model posits that attention improves population representation primarily by shaping neural tuning of individual units. Early studies using single-unit recordings in macaques showed that attention modifies the tuning curves of neurons in V4 for orientation^12,13^ and in the middle temporal area for motion direction^14^. In addition, attention has been shown to influence the spatial tuning of neurons in early visual areas^15,16^, with similar findings recently replicated at the level of single voxels in human imaging studies^17-20^. Furthermore, a substantial body of research has also examined how attention affects neuronal contrast response functions in macaques^21,22^ and humans^23,24^, inspiring several theoretical models^25,26^. These studies converge on the assumption that changes in neural tuning are the core mechanisms by which attention enhances population representations.

A competing view has recently emerged with the advent of multi-unit recording techniques. An increasing number of electrophysiological studies in macaques^8,9^,27 and neuroimaging studies^28^ in humans have suggested that the primary contribution to improved population representations may instead arise from changes in response correlations between units rather than from tuning modulations on single units. This “correlation change” model has gained supported in other cognitive domains, such as perceptual learning^29,30^, wakefulness^31^, and task engagement^32^. The ongoing debate between the tuning-based and correlation-based accounts underscores the importance of clarifying how attention shapes individual responses and their intrinsic relationship to population representations.

How can we resolve these competing models? We propose a framework of *neural population manifold analysis*^33-35^ that allow us to compare the effects of attention on tuning changes and correlation changes, and quantify their contributions to the overall fidelity of the population representation. This framework is valuable because characterizing neural population manifolds serves as an intermediate step to connect individual-level changes with improved population representations induced by attention. Therefore, this framework provides interpretable and falsifiable predictions that can be directly tested in empirically measured multivariate cortical responses. Addressing these questions is essential not only for understanding top-down modulation — such as how attention shapes visual processing — but also for elucidating the fundamental principles of information coding in the brain.

In this study, we combined fMRI, voxel-encoding modeling, and neural population manifold analysis to systematically investigate how attention improves population representations of spatial positions in the human visual cortex. Unlike previous fMRI studies that focused solely on measuring voxel tuning, we estimated both the tuning and covariance from thousands of trials of cortical responses while observers directed attention either to central fixation or to different stimulus positions across the visual field. Leveraging voxel-encoding modeling and multivariate decoding, we replicated many key findings of attention effects at both the individual level (e.g., changed tuning and reduced noise correlations) and population level (e.g., enhanced decoding accuracy). Most importantly, we conceptualized response changes of individual units into different types of population manifold transformations in a high-dimensional neural space. Our results reveal that tuning changes consistently enhance population representations across visual regions, whereas changes in variance and/or correlation changes have minimal—or even detrimental— effects. These findings resolve the long-standing debate and strongly support the “tuning change” theory of attention.

## RESULTS

In this study, we primarily focused on how cortical responses encode spatial positions across visual space. Inspired by previous fMRI studies on attention^17-19^, we presented a sequence of face stimuli at one position of a 4 × 4 spatial grid on each trial (Figure 1A). Human observers were instructed to perform either a one-back task on the digits at the central fixation (i.e., digit task), in which they attended exclusively to the central digits and ignored the faces, or a one-back task on the face identity (i.e., face task), in which they attended to the faces appearing at different locations while ignoring the digits. Importantly, in both tasks participants maintained central fixation, and the physical stimuli remained identical; the only difference was the attentional locus. We were primarily interested in how devoted attention to different spatial positions (i.e., face task), as compared to the attention to the central fixation (i.e., digit task), improves the overall spatial coding in human visual cortex. Note that our paradigm is different from the traditional posner cuing task where attention is directed a specific spatial location. Rather, the attention is directed to different stimulus position trial-by-trial. Importantly, unlike previous fMRI studies, we deliberately recorded many trials of cortical responses towards each position (9 sessions, 2560 trials in total for each subject), allowing us to systematically quantify both the tuning, trial-by-trial variance and correlation changes of voxels.

**Figure 1.**
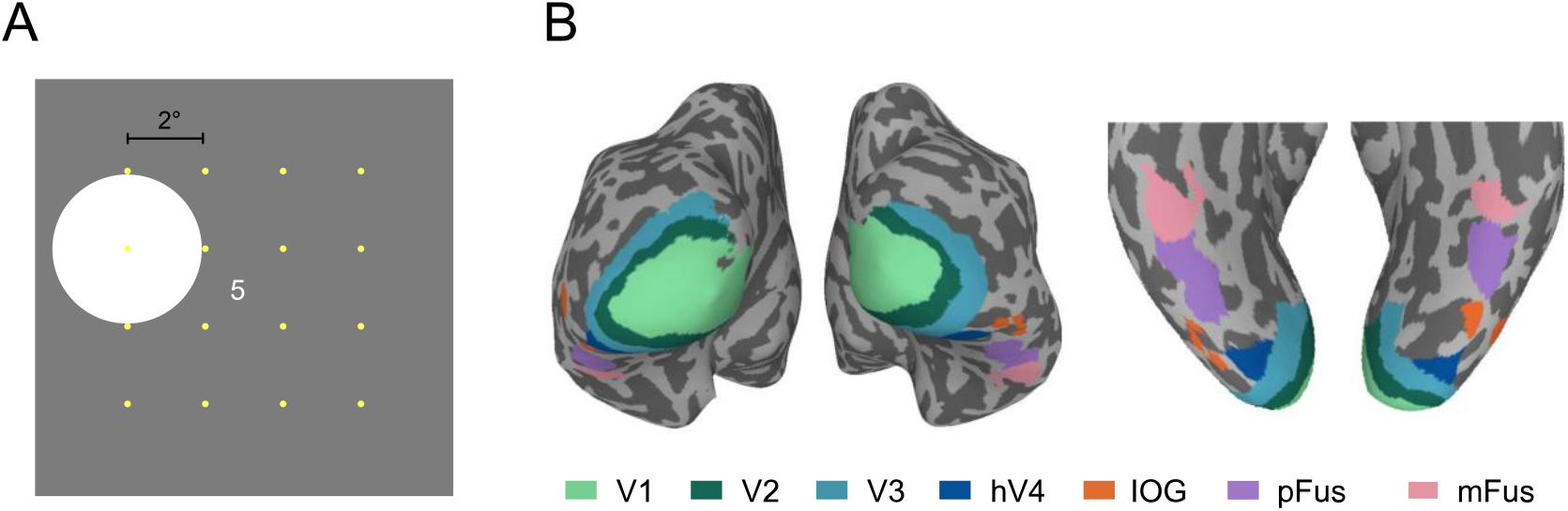
***A***. Experiment stimuli. A stream of digits was presented at the center of the screen, while a face appeared at one of 16 positions within 4 × 4 grid spaced 2° apart (position centers indicated as yellow dots, not shown in the actual experiment; the white circles indicate the positions where faces were presented). The face position switched systematically between trials. Within a trial, the face remained in the same location while its identity and viewpoint varied dynamically. In different runs, participants performed one of two tasks while maintaining central fixation: responding when the digit value repeated (digit task) or when the face identity repeated (face task). ***B***. ROIs for an example participant, where V1-hV4 were defined by a retinotopic experiment and IOG-mFus were defined by functional localizer experiments (see METHODS AND MATERIALS).

### Attention improves population representations in a distance-dependent manner

Previous studies have suggested that attention improves behavior by refining neural population representations of visual stimuli. We first examined whether attention indeed benefits population representations of spatial positions.

Consistent with decoding analysis in many fMRI studies, we trained a linear classifier (i.e., support vector machines) to decode any pair of spatial positions from multi-voxel responses. Given different spatial distances between position pairs, we divided our decoding analysis into two conditions: two positions either near (i.e., near-distance condition) or far (i.e., far-distance condition) from each other. We found that, compared to the digit task, the face task enhanced the position decoding accuracy in almost all regions (Figure 2A, Wilcoxon test, all *p*s < 0.001; except V1, *p* = 1) when the two positions were far apart. Such attention effects were less pronounced when the two positions were near. In the near-distance condition, the face task enhanced decoding accuracy only in hV4 and IOG (*ps* < 0.001) and even impaired decoding accuracy in mFus (*p* = 0.035). Such distance-dependent attention effects suggest that attention does not always benefit population representations for all stimulus conditions, even though the attention effects on individual responses are profound, highlighting the need to systematically quantify attention effects at both individual- (i.e., single voxels) and population-level (i.e., a pool of voxels).

**Figure 2.**
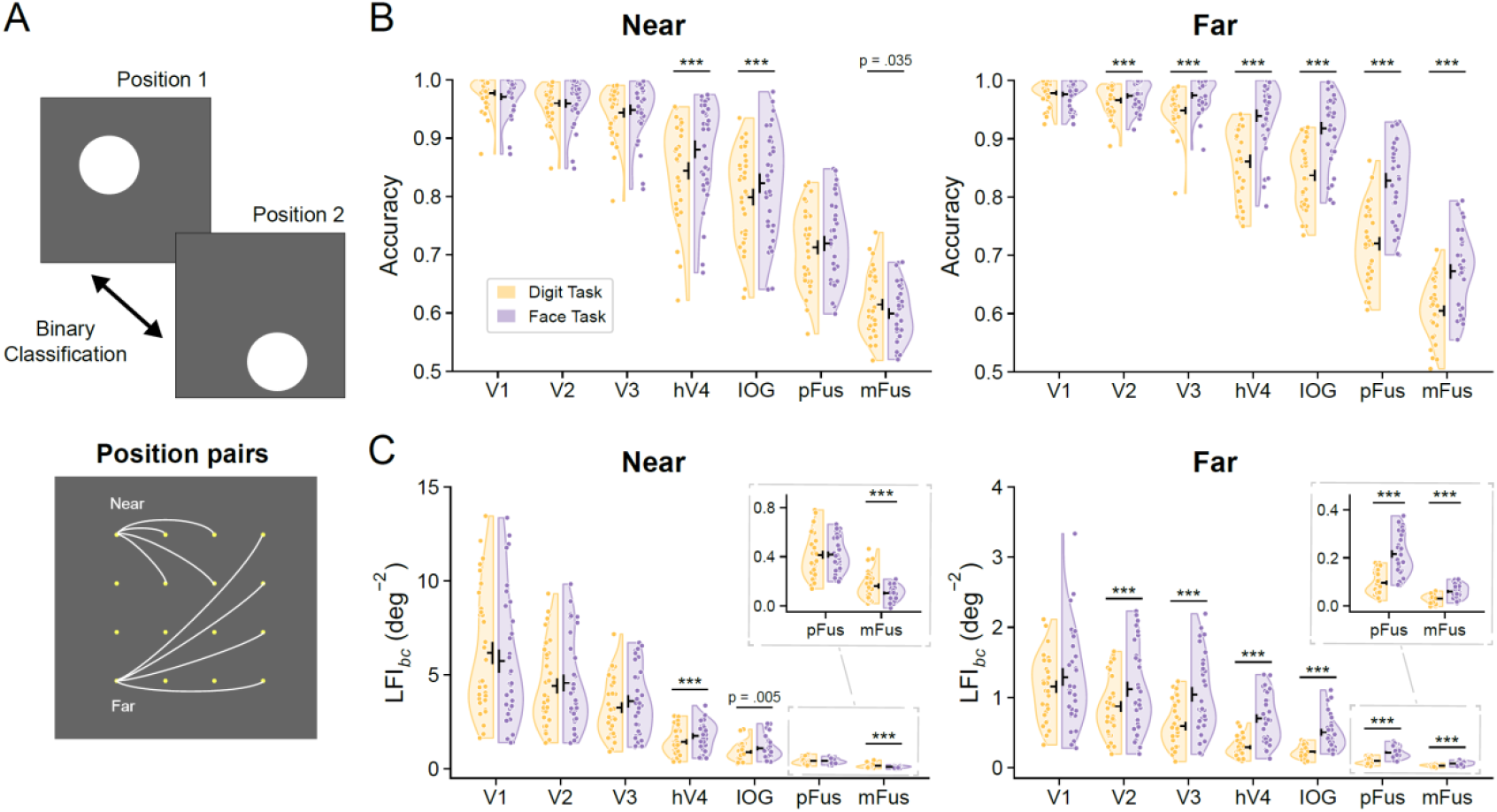
Attention effects on population representations of spatial positions. A binary classification was performed to decode any pair of positions based on multi-voxel responses in an ROI. All the position pairs resulted in 9 pairwise distances (2°, 2.83°, 4°, 4.47°, 5.66°, 6°, 6.32°, 7.21°, and 8.49°), binned into the near condition (first 4 distance bins, spatial distance < 4.5°) and the far condition (last 4 distance bins, spatial distance ≥ 6°). Linear SVM decoding accuracy (panel ***B***) and linear Fisher information (panel ***C***) of near (left) and far (right) position pairs for each task in each ROI. Each data point indicates a sample (8 participants × 4 distance bins = 32 samples). The black horizontal and vertical lines in the violin plots represent the mean and ± S.E.M. across samples, respectively. All significant differences are labeled, with three asterisks indicating *p* < 0.001.

We further introduced linear Fisher information—a standard metric to assess fidelity of population representation in computational neuroscience—to quantify the discriminability of two positions based on multi-voxel responses. The results echo the decoding analysis that the face task enhanced stimulus information in almost all regions (Figure 2B, Wilcoxon test, all *p*s < 0.001; except V1, *p* = 0.348) under the far-distance condition, and only enhanced stimulus information in hV4 (*p* < 0.001) and IOG (*p* = 0.005) under the near-distance condition.

Taken together, we found stimulus-specific attention effects on population representations in human visual cortex, and confirmed that attention indeed improves population representations of spatial positions, especially when two positions are far from each other.

### Attention changes spatial tuning in the human visual cortex

After validating the beneficial attention effects on population representations in our data, we next examined attention effects on response properties of individual units, consistent with previous approaches on single neurons in animal neurophysiological studies^15,16^,36,37 and single voxels in recent human imaging studies^18-20^.

We first examined the attention effects on the spatial tuning of individual voxels in human visual cortex. Throughout the paper, we called the receptive field of individual voxels as *voxel receptive field* (vRF) and used *population* to specifically indicate a pool of many voxels. We separately fitted the vRF model to the responses of each voxel and compared fitted vRF properties across the two tasks. Compared to the digit task, the face task slightly reduced the eccentricity of voxels in V1 (two-tailed sign test, *p* = 0.018) but significantly increased the eccentricity of voxels in later areas (V3-mFus, all *p*s < 0.001; Figure 3A). Similarly, the face task slightly reduced the vRF size of voxels in V1 and V2 (*p*s < 0.001), but significantly enlarged the vRF size in later areas (V3, IOG-mFus, all *p*s < 0.001; Figure 3B) except a non-significant effect in hV4 (*p* = 0.972). Moreover, the face task enhanced the amplitude of vRF in all ROIs (Figure 3C; *p*s < 0.001), suggesting that attention to stimuli generally enhanced voxel response amplitude along the human visual hierarchy. Interestingly, we found the attention effects on vRF tuning were generally stronger in high-level (e.g., pFus/mFus) than low-level visual areas (e.g., V1/V2). These results are qualitatively consistent with the previous fMRI study using the same paradigm^19^. In sum, our results here are consistent with the long-standing notion that attention can fundamentally shape neural tuning of individual units^17-19^.

**Figure 3.**
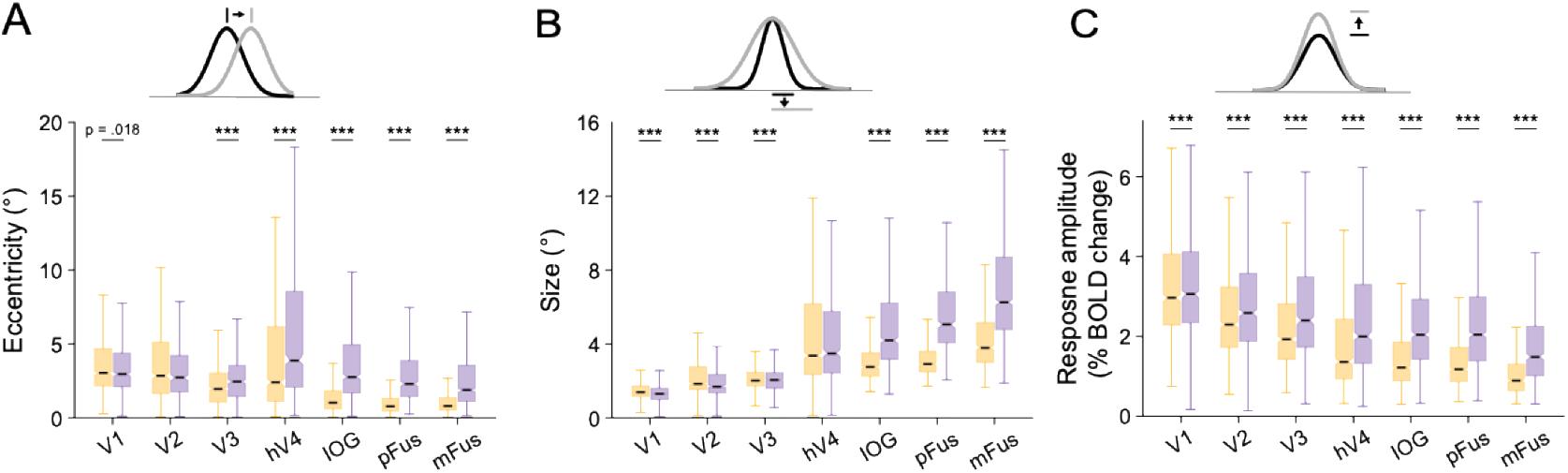
Attention modulation of vRF properties. Summary of eccentricity (panel ***A***), size (panel ***B***), and amplitude (panel ***C***) of vRFs in each ROI for each task. Here, vRF amplitude (i.e., scale factor) was empirically computed from the maximum predicted response amplitude rather than using the raw parameter. The black horizontal lines indicate median values across samples (100 voxels × 8 participants = 800 samples), notches indicate 95% CIs from 10,000 bootstraps, upper and lower hinges indicate 75th and 25th percentiles, respectively, and whiskers indicate 1.5× the interquartile range. All significant differences are labelled, with three asterisks indicating *p* < 0.001.

### Attention enhances response variance and reduces correlated activity in the human visual cortex

Previous fMRI studies in this line of research only focused on the attention effects on spatial tuning of individual voxels^17-19^. However, it is well-established in computational neuroscience that the fidelity of population representation depends on both tuning of units and covariance structure (i.e., response variance and correlations) between units^38,39^. The key advantage of our study is that we deliberately collected many trials of voxel responses towards each position, allowing us to systematically examine the covariance structure between voxels.

We first assessed the attention effects on the trial-by-trial response variance of single voxels in each position. We found that the face task enhanced voxel response variance in all ROIs (two-tailed sign test, all *p*s < 0.001; Figure 4A). Given that the face task also boosted the overall response amplitude of voxel responses (Figure 3A), we further examined the Fano factor (i.e., variance normalized to mean) of individual voxels. The results show that the Fano factor of voxel responses was reduced in all regions under the face task (two-tailed signed test, all *p*s < 0.001; Figure 4B). This finding is also consistent with the reduced Fano factor in monkey neurophysiological results^8,9^. Together, attention changes both voxel response amplitude and response variance, and attention improves the overall signal-to-noise ratio (i.e., inverse of Fano factor) of single voxels.

**Figure 4.**
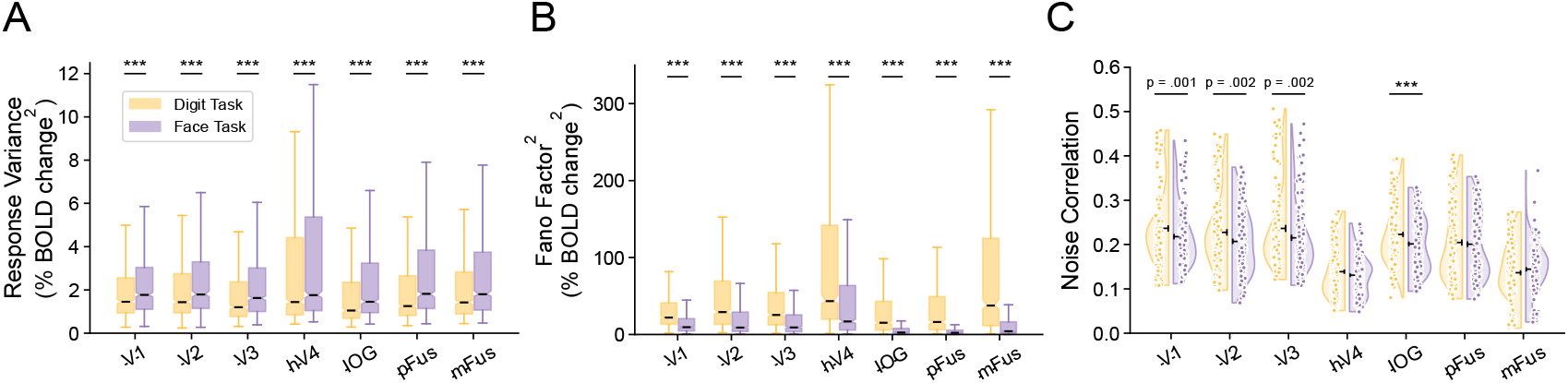
Attention effects on variance and correlations of voxel responses. The face task increases response variance (panel ***A***) and decreases the squared Fano factor (panel ***B***) in all ROIs. The black horizontal lines indicate median values across samples (100 voxels × 8 participants = 800 samples). The notches indicate 95% confidence intervals from 10,000 bootstraps. The upper and lower hinges indicate 75th and 25th percentiles, respectively. The whiskers indicate 1.5× the interquartile range. ***C***. Noise correlations for pairs of voxels in each ROI for each task. Each data point indicates a sample (8 participants × 16 positions = 128 samples). The black horizontal and vertical lines in the violin plot represent the mean and ± S.E.M. across samples, respectively. All significant differences are labelled, with three asterisks indicating *p* < 0.001.

Lastly, we examined the attention effects on the trial-by-trial response correlation (i.e., noise correlation) between voxels. We found that voxelwise noise correlations were mostly positive, but the magnitude was significantly reduced in the face task in several regions (V1-V3, IOG, all *p*s < 0.001; Figure 4C). This reduction of noise correlation by attention has also been documented in single neurons in previous electrophysiological studies^8,9^ and recently in an fMRI study^28^. This phenomenon is also thought to support the “correlation change” theory of attention. Previous monkey neurophysiological studies have also suggested that the changes in noise correlation between two units may depend on their tuning similarity^40^. We did not find such a relationship in our data (Supplementary Figure S1).

### Linking individual responses and population representations by neural population manifold

The above results suggest that attention fundamentally shapes various aspects of individual responses and improves population-level representations of spatial positions. Although several results reported here have been well documented in the literature, the key question that remains unaddressed is how changes in response properties of individual units contribute to improving population representations. The central debate lies in whether tuning changes or correlation changes serve as the predominant role of improved population representations.

Leveraging the approach of neural population manifold, we show in this section that response changes in individual units can be conceptualized into various forms of population manifold changes in high-dimensional neural space, which are naturally linked to changes in population representations. Given substantial attention enhancement of population representations in the far-distance conditions, we focused on the data in this condition (see Supplementary Figures S2-S3 and S6 for results in the near-distance conditions).

The neural population manifold approach assumes that trial-by-trial responses of *N* voxels towards two stimuli (i.e., two positions) form two neural manifolds (i.e., response distributions) in an *N*-dimension neural space. The tuning of individual voxels in a population together determines the mean values, and the variance and correlation of voxel responses (i.e., covariance) determine the shape of the two manifolds. The fidelity of population representations can be expressed by the discriminability of the two manifolds. Intuitively, there exist four potential mechanisms to improve manifold discriminability.

First, attention could enlarge the Euclidean distance between the mean values of the two distributions, a mechanism denoted here as *signal enhancement* (Figure 5A). As the mean values of the distributions are solely determined by individual tuning, signal enhancement corresponds to the “tuning change” theory of attention. Indeed, we found that the Euclidean distance (i.e., modulus of the vector) between the mean values of the two manifolds was significantly enhanced in the face task as compared to the digit task (Wilcoxon test, all *p*s < 0.001; Figure 5B).

**Figure 5.**
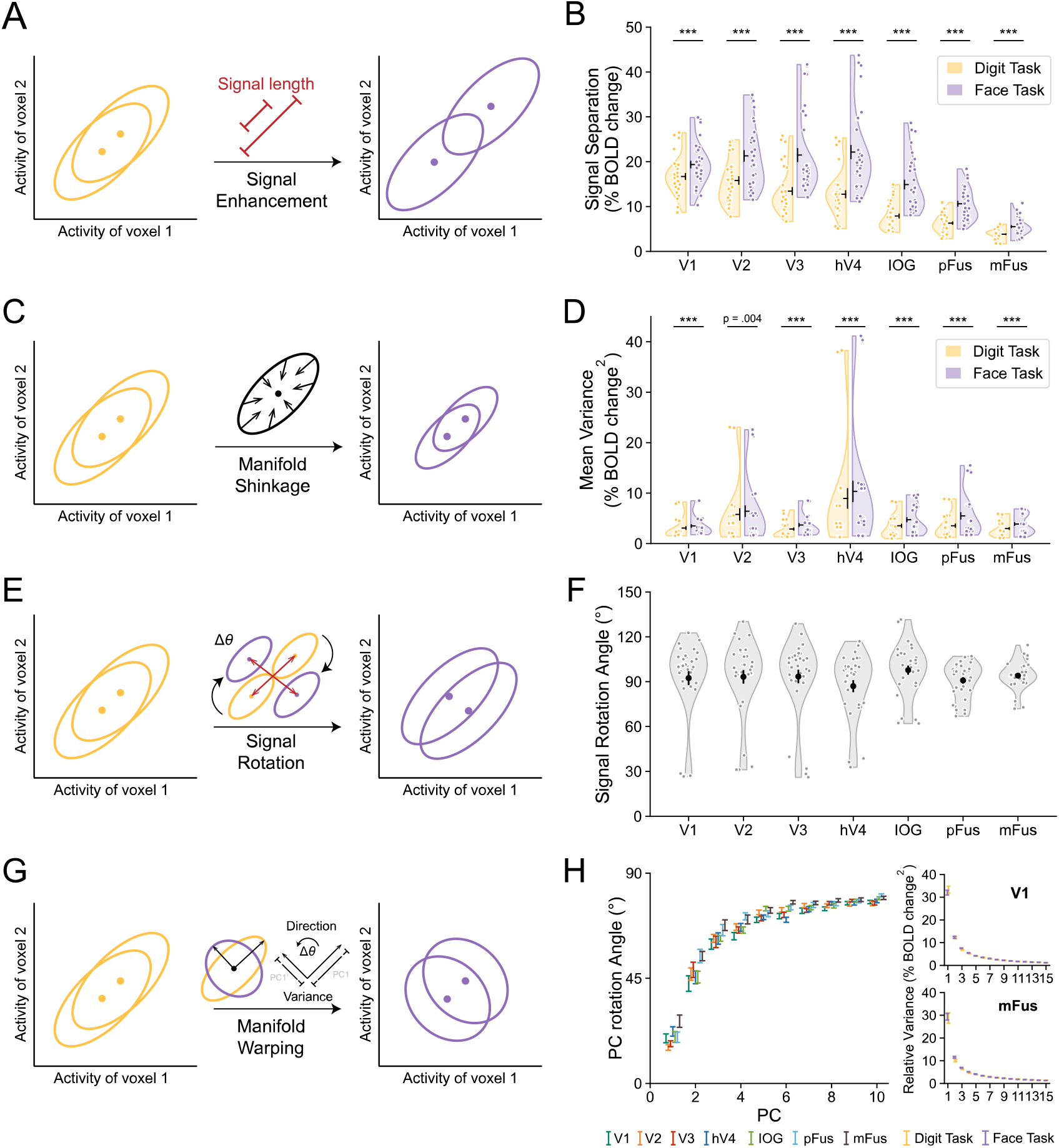
The theoretical illustration of the predictions of the four mechanisms (panels ***A/C****/****E****/****G***) and the empirical tests of these predictions (panels ***B****/****D****/****F****/****H***). In panel ***H***, eigen-decomposition was performed on the mean covariance. The rotation angles of the largest 10 PC from the digit task to the face task were plotted in all seven ROIs (left panel). The PC relative variance in both tasks in V1 and mFus (right panel). Each data point indicates a sample (8 participants × 4 distance bins = 32 samples). The mean ± S.E.M. were represented by black horizontal and vertical lines in (***B&D***), and by error bars in (***F&H***), respectively. All significant differences are labelled, with three asterisks indicating *p* < 0.001.

Second, attention could also shrink the overall size (i.e., response variance) of the manifolds, a mechanism denoted here as *manifold shrinkage* (Figure 5C). We calculated the overall response variance of the two distributions across two tasks. In contradiction to the prediction, the face task enhanced the overall voxel response variance rather than decreasing it (Wilcoxon test, all *p*s < 0.001; Figure 5D). The enhanced response variance in theory should impair population representation. This finding also highlights the complexity of this problem: not all changes at the individual level benefit population representation.

Third, attention could improve population representation by rotating their relative positions without changing the Euclidean distance between the mean values and the shape of the two manifolds (Figure 5E). This *signal rotation* mechanism predicts that the task rotates the two manifolds by a certain angle. Note that signal rotation also corresponds to the “tuning change” theory of attention. Indeed, we found that the direction of the signal vector rotated from the fixation to the face task, with the angles spanning a broad range (∼30º -130º) across all regions (Figure 5F).

Lastly, the *manifold warping* mechanism proposed that attention could improve population representation by changing the covariance structure of voxel responses (Figure 5G). This mechanism predicts that (1) the directions of the principal components of covariance changed across the two attention tasks, and (2) the variance redistributed across principal components. Indeed, we performed principal component analysis (PCA) and confirmed the two predictions (Figure 5H, V1 and mFus; Supplementary Figure S2F, other regions). This manifold warping mechanism is associated with the “correlation change” theory of attention.

In sum, by leveraging the neural population manifold approach, we can intuitively delineate the four potential manifold changes through which attention improves population representations. Importantly, the neural population manifold approach conceptualizes various forms of response changes at the individual level into interpretable geometric transformations. This approach also generates experimentally falsifiable predictions of these transformations. Note that by far, we only provide evidence for the existence of the four manifold changes in our empirical data. However, we emphasize that the existence of these ingredients does not guarantee that they pose *positive* effects on population representation, because their effects may interact and jointly determine population representation, as we will show in the next section.

### Quantifying the effects of manifold changes on population representation

Based on the qualitative illustration of the four possible manifold changes and their supporting evidence, we aimed to further leverage LFI to quantify the contribution of the four mechanisms.

We rewrote the formula of linear Fisher information, which represents the discriminability of two neural manifolds, as (see the proof in Supplementary Note 1)

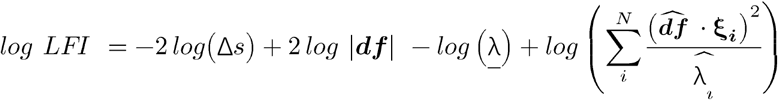

This equation provides several unique insights into the fidelity of population representations. Obviously, First, it suggests that the four mechanisms proposed above are logically complete— namely, attention improves population representations by these four and only four possible mechanisms. Specifically, signal separation (i.e.,|***df***|), mean variance (i.e., <u>*λ*</u>), the direction of the signal vector (i.e.,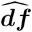), and covariance structure (i.e., ***ξ*** _*i*_ and *λ*_*i*_) all affects manifold discriminability. Second, this equation suggests that signal enhancement (i.e., increased |***df***|) and manifold shrinkage (i.e., reduced *λ*) independently enhance overall LFI (Figure 6A). But signal rotation and manifold warping interact, and they, as a unity, jointly contribute to LFI. In sum, our analytical derivation suggests that LFI is determined by three independent and linearly additive components. Our analysis here also summarizes the formal mathematical meaning of the four mechanisms and their corresponding geometric changes of neural manifolds.

**Figure 6.**
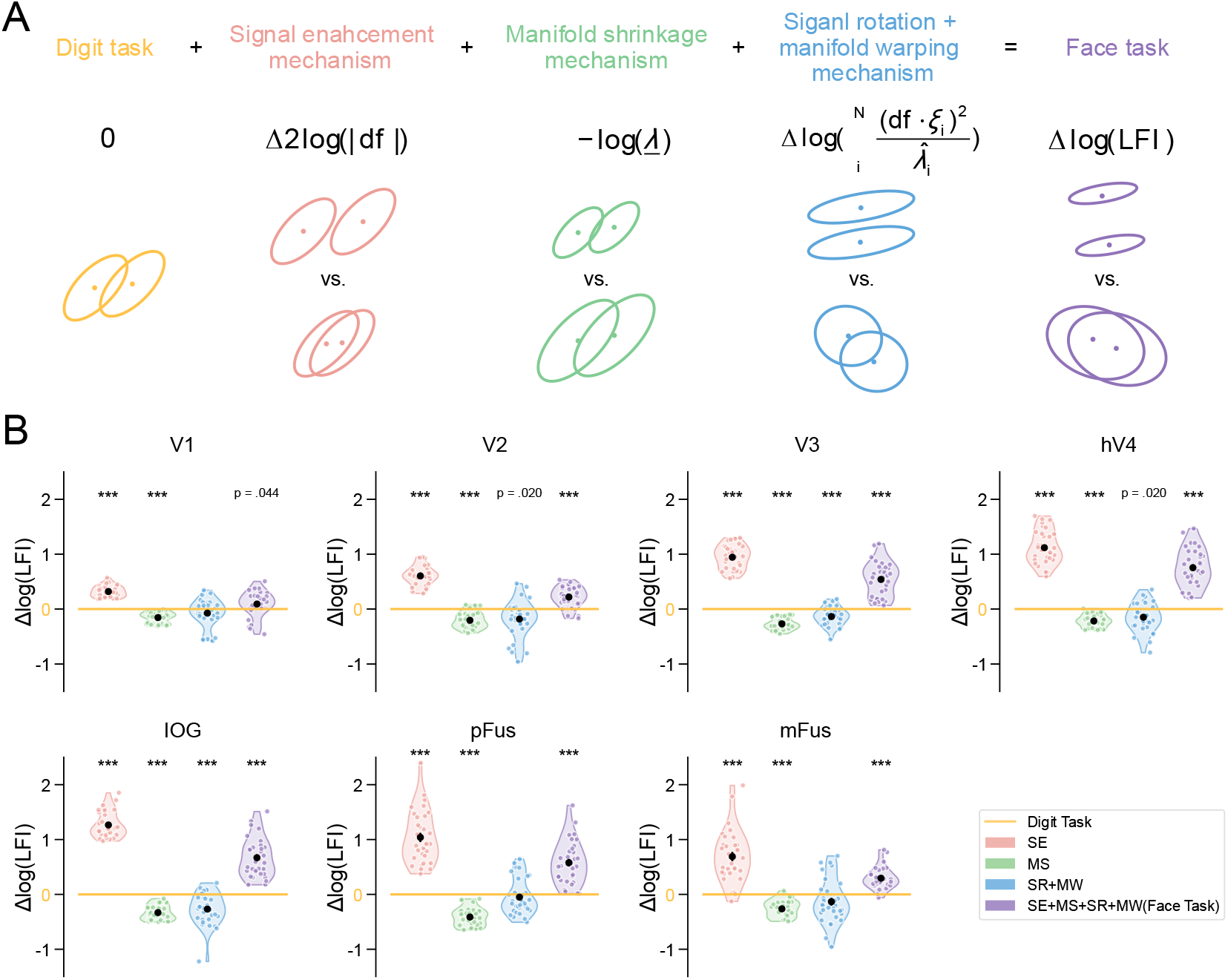
***A***. According to Equation 6, we tested the specific role of each mechanism on population representations in the face task separately. For each mechanism, we calculate the difference between the corresponding parameter in the LFI under the face task and the digit task, and examine whether attention effects on this parameter are beneficial or detrimental to population representations. Two of the mechanisms, signal rotation and manifold warping, were examined together because their interaction (blue data points) contributes to the LFI formula. ***B***. The difference between *log LFI* obtained by applying each mechanism or all mechanisms (i.e., the face task) and the *log LFI* of the digit task in each ROI. Each data point indicates a sample (8 participants × 4 distance bins = 32 samples). The error bars in the violin plot represent the mean ± S.E.M. across samples. All significant differences are labelled, with three asterisks indicating *p* < 0.001.

Our derivation also indicates important difference between our approach and the approaches in previous studies ^8,9^,40. We explicitly clarify that tuning changes produce two possible effects on signal vectors connecting the mean of two manifolds: enlarging the length of the signal vector (i.e., signal enhancement), which is independent of correlations, and changing its direction (i.e., signal rotation), which interacts with covariance. In other words, our framework indicates that tuning and correlation are naturally related, and thus it is impossible to evaluate the effects of correlation changes independent of tuning (Supplementary Figure S4).

By decomposing LFI into three independent and computable components, we are now able to evaluate their effects on population representation by directly calculating their changes from the digit task to the face task (Figure 6B). Consistent in all ROIs, the mechanism of signal enhancement induced by tuning changes is always beneficial to population representation. In contrast, we observed that enhanced response variance (i.e., manifold expansion) is always detrimental to population representation. Most importantly, the interaction of signal rotation and manifold warping shows little (i.e., V1/pFus/mFus) or even detrimental effects (i.e., V2/V3/hV4/IOG) to population representation. Notably, the net effects of attention on population representation, as the sum of the three components, are still positive. In other words, the net positive effects of attention on population representations are primarily dominated by signal enhancement, which is only induced by tuning changes of individual voxels. These results clearly suggest that not all the changes at the individual level benefit population representation. Our results strongly support the “tuning change” theory of visual attention and help resolve the long-standing debate on the neural mechanisms of attention.

### Attention effects manifest on small variance principal components

As the “correlation change” model has long been hypothesized as strong determinant of attention effects, the minimal or even detrimental effects of the term of tuning-correlation interaction is surprising. According to the Equation 5, the interaction between the tuning and covariance can be calculated as the sum of information across all principal components 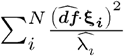. The information on each principal component is defined as the squared length of the projection from the unit signal vector 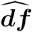 to the principal vector divided by the relative variance 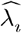 on that component. To better understand this term, we further deconstructed the interaction term 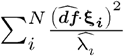 by examining the numerator 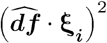, denominator 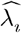, their division 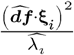, and the cumulated information of the first M PCs 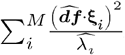 We especially focused on representative low-level visual area V2 (note the overall attention effects are not significant in V1) and high-level visual area mFus, where the contribution of the interaction terms is negative (see Supplementary Figure S5 for other regions).

Interestingly, we found both the numerator 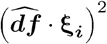 and denominator 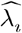 decreased as the PC rank increased (Figure 7A/B/E/F). Surprisingly, the information on a PC (their division) increased as the PC rank increased (Figure 7C&G). This effect suggests that the small variance PC provides stronger information to discriminate two stimuli, and the decreasing variance on each PC plays the predominant role of determining the role in each PC. Most importantly, the task effects on this term started to manifest from relatively small variance PC (∼20^th^ PC in V2, Figure 7C; ∼40^th^ PC in mFus, Figure 7G). The predominant effects of small variance PCs are more pronounced if we plotted the cumulated information across PCs (Figure 7D&H). In other words, the detrimental task effects mainly manifested on a small variance PC rather than the first large PCs. This is astonishing because this detrimental task effect will be completely overlooked if small variance PCs are discarded during the traditional principal component analysis. This effect also invites a rethinking the efficacy of PCA. The assumption of PCA is that covariance contains the useful information about a high-dimensional random variable, and that we can preserve the majority of information by only extract first a few principal components. Note that, here the objective is to mostly discriminate two sets of random variables. According to the definition of information of each PC 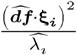, the small variance the larger information on that PC, which is fundamentally contradictory to the logic of PCA. Our results here challenge the long-standing convention of PCA when analyzing neural population manifold, discarding small-variable PCs in dimensionality reduction may overlook important aspects of information coding in neural populations.

**Figure 7.**
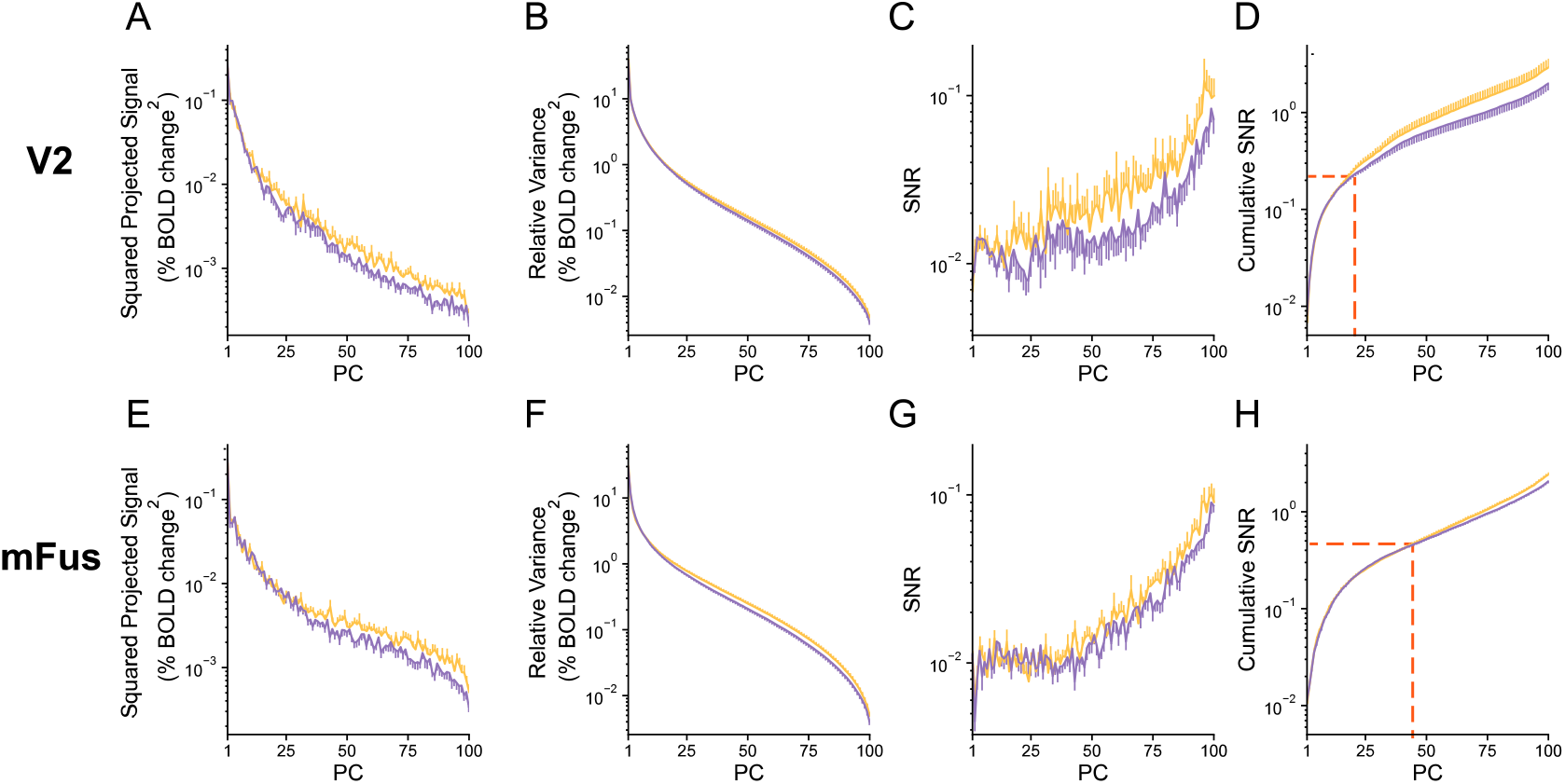
Eigen-decomposition analyses for the joint effects of signal rotation and manifold warping in V2 (***A-D***) and mFus (***E-H***). The x-axes are PCs ranked by eigenvalues from high to low. Each figure from left to right represented the squared projected signal 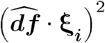 on each PC, the relative variance 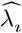 on each PC, the signal-to-noise ratio 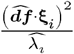 on each PC, and cumulative signal-to-noise ratio 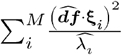 of the first *M* PCs. The mean ± S.E.M. across samples (8 participants × 4 distance bins = 32 samples) were represented by error bars, where only half of the error bars were drawn for better presentation.

## DISCUSSION

In this study, we investigated how attention improves population representations of spatial position in human visual cortex. Our results replicated a broad range of classical findings of attention at both the individual and population level. Leveraging the neural population manifold approach and linear Fisher information, we systematically quantified the effects of manifold changes associated with individual response changes. Surprisingly, we found that the manifold change induced by tuning changes accounts for almost all beneficial attention effects on population representations, while covariance (including correlation) changes consistently counteract the effects of tuning changes, resulting in minimal or even detrimental net effects. These results strongly support the “tuning change” theory of attention.

The primary goal of our study is to establish a connection between neural changes at the individual level and enhanced sensory representation at the population level. Here we summarize the overall logic flow of our inferences in order to clarify several important concepts in our study. First, in addition to the two well-documented factors—tuning and correlation changes—we show that the response variance of units is the third contributing factor because variance and correlation jointly determine covariance of population responses. Second, we conceptualize attention effects on tuning, correlation, and variance into four forms of geometric manifold changes: signal enhancement, manifold shrinkage, signal rotation, and manifold warping, and examined their existence in the multivariate cortical responses in our experiments. Third, based on the linear Fisher information, we analytically show that the four manifold changes can be further decomposed into three linearly additive terms: signal separation changes, manifold size changes, and rotation-covariance interaction. Finally, we quantify the three terms and find that the dominant role of signal enhancement in attention effects is rather than the other two terms.

Detailed quantifications of attention-related changes at both the individual and population level are of particular value to understanding information coding in the brain. A notable limitation of previous studies is that they usually only focus on a single aspect of response changes of individual units and without formal quantifications of their potential impact on population representations. For example, although many fMRI studies have shown that attention modulates spatial preferences of single voxels^17-19^, there was only one study attempting to link vRF changes to population coding of visual space^20^. However, that study did not take into consideration the effects of covariance of population responses. Here, our results employed a similar paradigm and demonstrate that significant vRF changes of single voxels do not necessarily improve or even worsen population coding of adjacent positions in some regions (e.g., mFus in Figure S2A). Likewise, we found that, while the phenomenon of reduced noise correlations has been observed in several cognitive processes, such as attention^8,9^,28, perceptual learning^29,30^,41, and deficits^42^, the phenomenon of reduced noise correlations *per se* provide no benefits for population representations in our study. Previous theoretical studies have prompted that what really matters here is the reduction of the specific form of correlations that limit information (e.g., differential correlations) rather than the general reduction of noise correlations^43^. In some instances, even an increase in noise correlations has proven beneficial^40^. In conclusion, the complex interaction between tuning and covariance suggests that the contribution to population representation should be considered when we evaluate attention effects on a specific property of individual responses.

Our study provides novel insights into several existing findings in this line of research^8,11^,17,20,40,41,44. Cohen et al.^8^ compared multi-unit activity in macaque V4 across different attention states, and also examined the contributions of tuning, variance, and correlations to population representation by projecting population activity onto a decision axis to obtain d-prime^33^. They concluded that changes in noise correlations accounted for nearly 80% of the attention effects on population representations. Their conclusions diverge from ours for two possible reasons. First, their analysis manipulated only one property (e.g., correlation) while holding others (e.g., tuning and variance) constant across attention states. As we have discussed above, this approach overlooks the interaction between tuning and covariance and the fact that attention changes all response properties simultaneously. However, both empirical analysis and analytical derivation suggest that, while manipulating tuning or correlation only may lead to notable changes in population representation, the interaction between these factors could nullify their individual effects. Recently, we observed this exact complex interaction in our recent work on visual perceptual learning^33^. Therefore, it is impossible to evaluate the effects of correlations without considering tuning and variance changes. Second, the discrepancy may also reflect the intrinsic differences between voxel activity in fMRI and neuronal activity in neurophysiology. Specifically, fMRI voxel activity represents the aggregated responses of many underlying neurons within a voxel. We have previously shown that the distinction between voxel and neuronal activity may lead to differing effects of noise correlation^34^. We emphasize that the neural manifold approach is a unified computational framework to study multivariate responses measured by different modalities, including multi-unit recording data. Future studies should explore the contributions of these mechanisms across a range of neural measurement techniques.

Our study emphasizes the comparison of individual and population-level changes across different attention states, which distinguishes it from several theoretical work that merely compares population representations with and without noise correlations. For instance, many theoretical and empirical studies compared population responses in the case of all noise correlation being intact and the case of shuffling multi-trial population responses to disrupt noise correlations completely^28,34^,45,46. In such analyses, noise correlations are typically found to be detrimental, leading to the conclusion that reducing noise correlations should improve population representations^28,34^,45. Using a similar shuffling approach, we examined this effect and replicated the findings of a previous fMRI study^28^ on attention (Supplementary Figure S7). We emphasize, however, that this type of analysis (i.e., with vs. without noise correlations) fundamentally differs from the analysis of population representation changes under different cognitive states (i.e., attention vs. unattended). In both attention tasks, noise correlations are present in the data, yet their structure varies substantially with the attention state. Since attention modulates not only correlations but also tuning and other response properties, it is insufficient to infer the effects of noise correlation changes solely by manipulating correlation while keeping other factors constant.

Our finding that the detrimental effects of tuning-correlation interaction mainly manifest in small variance PCs challenges the long convention of using PCA for dimensionality reduction when analyzing population activity. The basic assumption of PCA is that population responses lie in the low-dimensional space due to intrinsic correlations between neurons, and we can reduce the dimensionality by only preserving several large variance PCs. However, this assumption may not hold in the case of discrimination tasks. The term 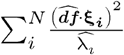 clearly demonstrates that, given a constant numerator, the smaller variance (i.e.,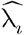) in a PC the more information (i.e., usefulness to discriminate two stimuli) a PC contains. In this case, we have to rethink the efficacy of PCA. We argue that analyzing data by eigen-decomposition is useful, but discarding small variance PCs should be careful. In general, we argue that estimating covariance is as important as measuring neural tuning in understanding population coding, which is usually ignored due to limited number of trials in previous fMRI studies.

Our studies also face several limitations. First, here we only focus on how attention improves population encoding of a specific visual feature (i.e., spatial position in our case)— namely, attention enhances the maximum amount of information an optimal decoder can possibly read out from population activity. But it remains unclear whether attention improves behavior by also optimizing the readout decoder. Some recent evidence suggested that optimizing the readout may also act as an important way to improve behaviour^47-51^. Future studies are needed to systematically disentangle the effects of attention on information encoding and decoding. Second, in order to optimize our fMRI experimental design, we asked participants to perform a one-back task on face identity. However, the visual feature we decoded is spatial position, which is in theory task-irrelevant. We did not used the traditional Posner cuing paradigm as the cuing paradigm requires a larger number of trials for cued (e.g., 70% trials) as compared uncued trials (e.g., 30% trials), which is suboptimal for decoding analysis. Consistent with several classical attention experiments in human imaging and animal neurophysiology^16,17^,20,21,36,37,52-54, the only purpose of the face task is to keep participants’ attention on the stimuli. Thus, we would treat our comparison as “attention to stimuli” vs. “attention to fixation” rather than the traditional definition of spatial attention where attention should be deployed to two particular positions. Third, although our fMRI results resemble many classical findings in both monkey neurophysiology and human imaging, future studies using large-scale in vivo population recording in animals are needed because fMRI and unit-recording studies measured intrinsically different cortical responses at different spatial scales. More sophisticated in vivo multi-unit recording techniques may shed new light on the nature of population codes in the brain.

## METHODS AND MATERIALS

### Ethics statement and participants

Eight volunteers (23-25 years old, 3 males and 5 females) with normal or corrected-to-normal vision participated in this study, including one of the authors (S.L.). The experimental protocols were approved by the Research Participants Review Board at Shanghai Jiao Tong University. All participants provided a signed informed consent form.

### Stimuli and Experimental Settings

#### Apparatus

Visual stimuli were generated by a Windows PC using MATLAB Psychtoolbox^55^ and presented by a Digital Light Processing System positioned at the head of the scanner bed. The monitor operated at a refresh rate of 60 Hz and a resolution of 1024 × 768 pixels in the retinotopic mapping experiment and 1920 × 1080 pixels in other experiments. Participants viewed the visual display through an angled mirror mounted on the head coil, with a field of view of 31° × 18° (viewing distance = 162 cm). Behavioral responses were recorded using a button box.

#### Main attention experiment

The fMRI experiments were adapted from Kay, et al. ^19^. Throughout the whole experiment, participants maintained a central fixation on a string of digits while face images were presented at one of 16 positions on a gray screen (Figure 1A). Participants were instructed to complete either a digit or a face one-back task while observing the stimuli in each run. During the digit task, participants were asked to maintain fixation on the central digits and to press a button when the digit repeated the value of the previous digit (i.e., one-back digit task). In the digit task, participants attend exclusively to the digits while completely ignoring the face images. During the face task, participants were asked to maintain fixation on the central digit and to press a button when a face identity was consistent with the previous face (i.e., one-back face task). In the face task, participants were required to attend exclusively to the face stimulus while completely ignoring the digits. We emphasize that the physical stimuli presented in both tasks were identical, and the only difference was whether attention was directed to the central digits in the digit task or to faces appearing in different positions in the face task.

Throughout the experiment, a 0.6° × 0.6° digit (varying between 0 and 9) was presented consistently at the center of the screen. The digit value (0-9) changed every 0.5 s. Digit repetitions occurred with a probability of 0.088, with a maximum of two consecutive repetitions allowed. To attenuate visual brightness adaptation, the digit color was alternated between black and white on consecutive presentations. On each valid trial, a sequence of four faces was presented at one of 16 positions with a 4 × 4 grid on the screen. The 4 × 4 position grid was spaced at 2° apart, and the radius of the circular face image was 2°. The face images were derived from the stimuli set by Kay, et al. ^19^, in which circular images were generated for each individual at 7 viewpoints (0°, ±15°, ±30°, ±45°) and converted to grayscale. The positions of the four faces were identical, but the individual identities and viewpoints differed. Each sequence had a 0.9 probability of containing consecutive repetitions of the face identity, allowing up to three consecutive repetitions. Each face sequence consisted of four different face images, each of which was presented sequentially for 1 s. Each valid trial consisted of a 4-s sequence of faces followed by a 1-s blank. Note that the overall frequency of digit repetitions was roughly matched to the overall frequency of face-identity repetitions.

Each scanning run consisted of an initial fixation period (8 s), 40 trials (5 s per trial), and a final fixation period (8 s), for a total duration of 216 s. The 40 trials were composed of 32 valid trials (16 position conditions, each presented twice in a randomized order) and 8 blank trials (no face stimuli presented, randomly interspersed). Each session consisted of 10 runs, with the digit and face tasks alternating for 5 runs each. To ensure that the only difference between the two tasks was the participant’s attention state, we pseudo-randomized five generated sequences of digits and faces between runs of each task, so that the stimulus sequences seen by participants in the two tasks were consistent. To collect sufficient trials in each condition, each participant completed 8 valid scanning sessions of the main experiment over multiple days. Thus, each participant completed a total of 2 tasks × 16 positions × 2 trials × 5 runs × 8 sessions = 2560 valid trials. To check whether participants were fixating accurately at the center of the screen throughout the experiment, we recorded their left eye movements at a sampling rate of 500 Hz using a scanner-compatible Eyelink 1000 eye tracker (SR Research).

#### Retinotopic mapping experiment

The procedures for retinotopic mapping experiment were consistent with those reported in Benson, et al. ^56^. Visual stimuli consisted of slowly moving apertures and dynamic, colored textures placed within the apertures. The stimuli were generated at a resolution of 768 × 768 pixels and were presented on a gray background. The experiment consisted of four runs, each lasting 300 s. Two types of aperture sequences alternated twice, so that the order of the runs was: MULTIBAR, WEDGE-RING, MULTIBAR, and WEDGE-RING. The MULTIBAR sequence comprised bars sweeping in eight different directions. The bar swept for 28 s in each direction, followed by a 4 s pause. Each run comprised a 16-s blank period, four 32-s sweeping blocks (right, up, left, down), a 12-s blank period, four 32-s sweeping blocks (upper-right, upper-left, lower-left, lower-right), and a 16-s blank period. The WEDGE-RING sequence comprised rotating wedges and expanding and contracting rings. The rotating wedges rotated in a clockwise or counterclockwise direction in each 32-s block. The rings expanded outward or contracted toward the center for 28 s and then rested for 4 s in each block. Each run comprised a 22-s blank period, eight 32-s sweeping blocks (counterclockwise rotating wedges, counterclockwise rotating wedges, expanding rings, expanding rings, clockwise rotating wedges, clockwise rotating wedges, contracting rings, contracting rings), and a 22-s blank period. The colored textures within the apertures comprised colored objects of multiple scales placed on an achromatic pink noise background. The apertures and textures were refreshed at 15 Hz. In the experiments, a 0.23° × 0.23° dot at the center of the display randomly changed to one of three colors (black, white, or red) every 1-5 s. Participants were asked to maintain fixation on the central dot and to press a button when the dot color changed. To enhance the stability of central fixation, a semi-transparent grid was added to the visual display^57^.

#### Functional localizer experiments

The functional localizer (fLoc) experiment, developed by Stigliani et al.^58^, was used to identify face-selective regions. Visual stimuli were images (1024 × 1024 pixels) comprising items overlaid on the phase-scrambled background, presented on a gray background. The items were classified into five different domains, with each domain comprising two categories: characters (numbers or words), bodies (with obscured heads or limbs), faces (adults or children), places (corridors or houses), and objects (cars or instruments). A mini-block design was used in the experiment. Each 4-s block comprised 8 stimuli of the same categories, with each stimulus presented for 0.5 s. For each experimental run, the order of the five stimulus domains and the blank baseline was counterbalanced, resulting in a total of 6 conditions. Each condition was presented in 12 blocks, with 6 blocks for each category within a stimulus domain. Each run consisted of a 12-s countdown, 72 4-s blocks, and a 12-s blank rest period, for a total duration of 312 s. Participants were asked to fixate on a central dot and to press a button when a phase-scrambled oddball picture appeared. Each run comprised 20 oddball images, with a maximum of one oddball image per block. The frequency of oddball images was equal across stimulus categories.

### MRI data acquisition

MRI data were acquired using a united-imaging 3T uMR 890 scanner with a 64-channel head coil. All the functional data were acquired from T2*-weighted gradient echo planar imaging (EPI) with 60 slices and a voxel size of 2.5 × 2.5 × 2.5 mm (TR = 2000 ms, TE = 30 ms, flip angle = 81°, Field of View = 200 × 200 mm; 2 × simultaneous-multi-slice acceleration; A→P phase encoding). In each scanning session, we interspersed three short EPI runs (20 volumes) with the slice prescriptions identical to the functional runs except for the reverse phase-encoded direction (P→A). These short EPI runs were used to correct for local spatial distortions. For each participant, we also collected a high-resolution T1-weighted MPRAGE anatomical scan (TR = 8.3 ms, TE = 2.2 ms, flip angle = 8°, Field of View = 256 × 256 mm, slices = 240, 1-mm isotropic voxels).

### MRI data preprocessing

Cortical reconstruction was performed on T1-weighted anatomical images using the standard procedures in FreeSurfer v7.3.2^59,60^. All the functional imaging data were preprocessed using customized scripts based on AFNI^61^. We first removed spikes from the time series and aligned all slices from each EPI brain volume to the same time point. Then, the functional data were reverse-polarity phase-encoding corrected, motion corrected (6-parameter affine transform), and aligned to individual anatomical images. For the retinotopic mapping and fLoc experiment, the data were additionally performed volume-to-surface mapping using the AFNI (SUMA) standard grid (std. 141)^62,63^ and spatial smoothing with a 4 m FWHM Gaussian kernel. For the main experiment, analyses were performed in the volume space and no spatial smoothing was applied. Finally, the BOLD time series of each voxel or vertex were scaled to have a mean of 100, yielding units of percent signal difference from the mean for each run.

### ROI definition

All ROIs were defined in the participants’ native anatomical volume space based on functional localizer experiments using the SUMA in AFNI. For the early visual area, we used the MATLAB toolbox analyzePRF (http://cvnlab.net/analyzePRF/) to fit a voxel receptive field (vRF) model with compressed spatial summation (CSS) to the BOLD time series of the retinotopic mapping runs for each vertex^64,65^. The preferred polar phase angle and eccentricity of vertices estimated by the vRF model were then visualized on the participants’ cortical surface. A threshold of greater than 10% response variability explained by the vRF model was set and V1, V1, V2, V3, and hV4 were then drawn by visual inspection.

For the face-selective areas, we used a general linear model (GLM) to fit the BOLD time series of fLoc runs and estimated the response amplitudes to different stimulus categories for each vertex. Based on the contrast map (face > text, body, place, object; *t* > 3, vertex level, uncorrected) and anatomical locations^66,67^, we defined the middle fusiform gyrus (mFus), the posterior fusiform gyrus (pFus), and the inferior occipital gyrus (IOG).

For each ROI, the vertices defined on the cortical surface were remapped to the native volume space to identify the corresponding voxels, which were used in the subsequent analyses for the main experiment.

### fMRI general linear modeling

We used the AFNI 3dDeconvolve function to fit a general linear model (GLM) to the BOLD time series of the main experiment. Specifically, we used the *-stim_times_IM* option of the function to include each valid trial (i.e., the 4-s trial containing a face sequence) as an independent predictor of the model to estimate the percent signal change from baseline for each voxel in each trial. The GLM also included demeaned head-motion parameters as well as constant, linear, and quadratic polynomial terms as nuisance predictors. For each participant, we concatenated the time series of all the runs during each scanning session and built a GLM. Subsequent analyses of the main experiment data were conducted based on the response amplitudes estimated by the GLMs (i.e., beta weights, in units of % BOLD signal change) at the voxel level.

### Voxel selection

To make the attention effects between the different ROIs quantitatively comparable, we selected 100 voxels within each ROI. Specifically, we combined voxels from both hemispheres to account for spatial representations of the full view-field. First, we took the mean of the GLM goodness-of-fit *R* ^2^ (see “fMRI general linear modeling”) across 8 sessions and selected the 100 voxels with the highest vRF goodness-of-fit *R* ^2^ in the digit task (see “Individual voxel receptive field analysis”). Consequently, the selected voxels exhibited a higher *R* ^2^ for both GLM and vRF fitting.

### Analysis of population representations

#### SVM decoding

To examine the effect of attention on the accuracy of spatial representations, we performed decoding analyses on population responses in two tasks. We trained linear support vector machines (SVM) using Python’s scikit-learn toolkit. The binary SVM classifiers were trained to decode any two different positions (position pairs: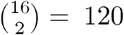) by the response amplitudes of 100 selected voxels in each ROI under two tasks. Response amplitudes were not normalized to preserve correlations, and 10-fold cross-validation was used.

To summarize the results, 16 spatial positions result in 120 position pairs, and 120 position pairs were divided into 9 bins based on the distance between them. The distances for 9 bins were 2°, 2.83°, 4°, 4.47°, 5.66°, 6°, 6.32°, 7.21°, and 8.49°, with each bin containing 24, 18, 16, 24, 8, 8, 12, 8, and 2 pairs, respectively. The first four distance bins were combined as the *near-distance* condition, and the last four distance bins were combined as the *far-distance* condition. The mean classification accuracy was calculated over the position pairs within each distance bin. For each task, each distance condition, and each ROI, the mean accuracy of each distance condition was pooled across distance bins and participants.

#### Linear Fisher information

To quantify the effect of attention on spatial representations by population codes, we calculated the linear Fisher information (LFI). LFI is a standard metric used to quantify the amount of information contained in observed neural responses relative to the stimulus variable. In this study, the linear Fisher information was used as a measure of the ability of the population responses to discriminate between two spatial positions.

On each trial, a stimulus elicited the responses of a population of voxels. In general, given *N* voxels in an ROI, the voxel population response in each trial can thus be viewed as a point in an *N*-dimensional neural response space, where each dimension corresponds to a single voxel. In each task, we collected *T* trials for each position. The population responses to each position form an *N* × *T* matrix, corresponding to an *N*-dimensional response distribution. Given any two positions *s*_1_ and *s*_2_, the LFI to discriminate the two *N*-dimensional response distributions is defined as:

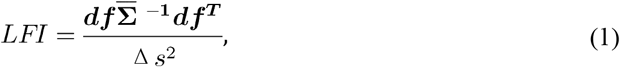

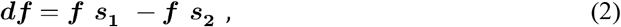

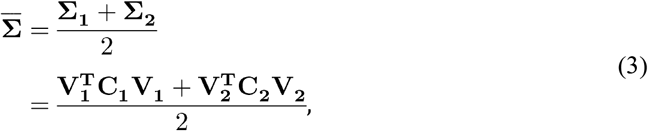

where ***f*** (***s***_**1**_) and ***f*** (***s***_**2**_) were *N*-dimensional response vectors, denoting the mean population responses to *s*_1_ and *s*_2_ across trials, i.e., the centers of the high-dimensional response distributions of *s*_1_and *s*_2_, respectively. ***df*** was the signal vector (ℝ^1×*N*^) connecting the centers of the two distributions, representing the difference between the two mean population responses. **∑**_**1**_ and **∑**_**2**_ denoted the covariance matrices (ℝ^*N*×*N*^) of the population responses to *s*_1_and *s*_2_, respectively. ***V*** was the diagonal matrix (ℝ^*N*×*N*^) with standard deviation, and ***C*** was the response correlation matrix 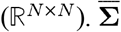 represented the mean of the two response covariance matrices. Δs denoted the physical distance (in visual degree units) between the position pair.

In practice, it is difficult to accurately estimate the signal vector and the covariance matrix from a finite number of trials. As a result, the estimate of linear Fisher information from Equation 1 can be substantially biased, due to its non-linear dependence on the estimated signal and covariance based on limited measurement^38^. Fortunately, it was previously shown that the bias can be analytically corrected^38^. The bias-corrected linear Fisher information, *LFI*_*bc*_, was be computed as follows:

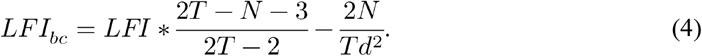

The bias-corrected method is only valid when *T* > (*N*+2)/2, which is satisfied in the present study (*N =* 100, *T* = 80). The mean bias-corrected LFI was calculated across position pairs within each distance bin. For each task, each distance condition, and each ROI, the mean bias-corrected LFI of was pooled across distance bins and participants.

### Analysis of individual voxel responses

To understand the effect of attention on spatial representations at the level of individual voxels, we quantified the voxel receptive field properties, Fano factor, and noise correlation in two tasks.

#### Individual voxel receptive field analysis

We fitted the voxel receptive field (vRF) for each voxel in the two tasks. To distinguish between the “population” of neurons within a voxel described here and the “population” of voxels described above, we referred to the receptive field of a voxel as vRF to emphasize its specificity to a single voxel, and used “population” to specifically indicate a pool of voxels. For each voxel, we fitted a vRF model with a compressive spatial summation (CSS) to the response amplitudes^64,65^ (http://kendrickkay.net/socmodel/) in each task. The CSS model predicts the responses of each voxel to the stimulus presented at any position in the visual field by computing a weighted sum of the contrast stimulus and an isotropic 2D Gaussian and then applying a static power-law nonlinearity. The model parameters include the center (*x, y*) and the size (*σ*) of the 2D Gaussian, the power-law exponent (*n*), and the overall amplitude (i.e., scale factor) of the predicted responses (*g*). For the purpose of modeling, the stimulus display was transformed into a square contrast image of 1080 × 1080 (considering only the face stimulus and ignoring the central digit) and then downsampled to 200 × 200. Negative response amplitudes were rectified to 0.

We quantified the vRF properties by defining the vRF eccentricity as 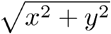, vRF size as 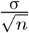, and vRF amplitude as the maximum predicted response to the stimulus in the model fitting (yielding an empirical measure that is more robust than the raw amplitude parameter^19^). The properties of selected voxels were pooled across participants in each task for each ROI.

#### Fano factor

The Fano factor is originally defined as the variance-to-mean ratio of neuronal spike counts in a time window. Here, we defined it analogously as the variance-to-mean ratio of voxel response amplitudes across trials. Considering the possibility of negative response values, we used the squared Fano factor. For each voxel, we calculated the squared Fano factor from responses in 80 repeated trials at each position and took the median across 16 positions. The squared Fano factors of selected voxels were pooled across participants in each task for each ROI.

#### Noise correlation

The noise correlation was calculated as the Pearson correlation coefficient of the responses to the 80 repeated trials of identical positions between two voxels. For each participant and each position, the noise correlation matrix was computed for 100 selected voxels within an ROI. We took the median of the lower triangle of the matrix and performed Fisher’s z-transformation for statistical tests. For each task and each ROI, the coefficients were pooled across 16 positions and 8 participants.

### Analytical derivation of LFI and four manifold changes

To further understand how attention influences population representations, we analyzed four potential modulation mechanisms.

In the LFI, we performed an eigen-decomposition on the mean covariance matrix 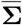 (see Supplementary Note 1 for detailed derivation), so Equation 1 can be rewritten as:

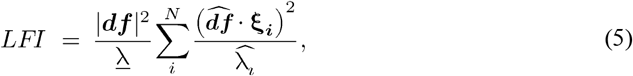

where | ***df*** |is the modulus length of the signal vector. <u>*λ*</u> denotes the mean of the eigenvalues.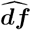 is the unit vector, representing the direction of the signal vector, where 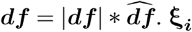 is the *i-*th eigenvector which denotes the direction of the *i-*th dimension. 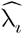 is the relative variance of the *i-*th dimension, where 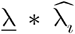 is the *i-*th eigenvalue.

According to Equation 5, we could infer four potential mechanisms of attention that affect the information encoded by the population responses: (1) signal enhancement, the increase in the distance between the mean responses to *s*_1_ and *s*_2_, as reflected by the signal separation | ***df*** |; (2) manifold shrinkage, the decrease in the size of the response distributions to *s*_1_ and *s*_2_, as reflected by the mean variance <u>*λ*</u>; (3) signal rotation, the rotation of the direction of the signal vector in favor of discrimination, as reflected by the unit signal vector 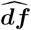; (4) manifold warping, the change in the shape of the response distributions (i.e., the direction and variance of each dimension) in favor of discrimination, as reflected by both *ξ*_*i*_ and 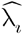.

To examine which mechanisms were involved, we first analyzed quantitative metrics for each of the four potential mechanisms. Signal separation, | ***df*** |, was computed as the modulus length of the signal vector for each position pair under each task. Mean variance, *λ*, was obtained by averaging the eigenvalues of the mean covariance for each position pair under each task. Signal rotation angle, *θ*^*s*^, was calculated as the angle between the unit signal vectors of the digit and face tasks for each position pair, with the range of [0, *π*). Covariance rotation angle,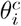, was calculated as the angle of the *i-*th eigenvector (i.e., *ξ* _*i*_) of the mean covariance for each position pair between the digit and face tasks, with the range of [0, *π*/ 2). Additionally, the relative variance of the covariance,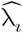, was computed for each position pair under each task. For further analysis and visualization, the signal separation and mean variance were averaged across position pairs within each distance bin. For each task, each distance condition, and each ROI, the mean was then pooled across the corresponding distance bins and participants. The signal rotation angle was averaged across position pairs within each distance bin and pooled across distance bins and participants for each distance condition and each ROI. The same operation was performed on the covariance rotation angle for each dimension and on the covariance relative variance for each dimension and each task.

To quantify the unique contribution of each attention mechanism, we proceeded to apply substitution to the LFI formula. Based on the population responses to the digit and face tasks, the parameters 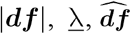, and ***ξ*** _*i*_ and 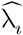 can be computed under each task. In this section, we described the replacement of parameters based on LFI under the digit task with those computed from the face task, and the subsequent comparison of the changes in LFI, to check the corresponding attentional modulation mechanism. The details were described as follows.

For convenience, we took the logarithm of the LFI:

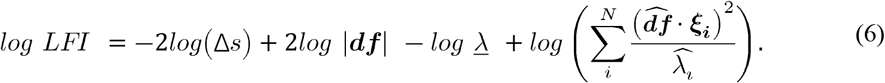

Thus, the information changes of population activity can be expressed as:

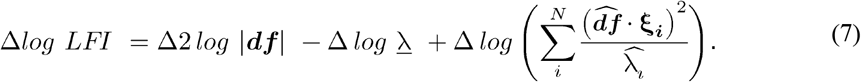

In other words, the task induced attention effects on population representations (Δ*ln LFI*)) can be decomposed into three linearly additive terms. We thus can directly calculate each term.

The contribution of signal enhancement (Δ *ln*(|***df***|)) and manifold shrinkage (− Δ *ln* (*λ*)) were quantified by:

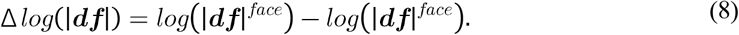

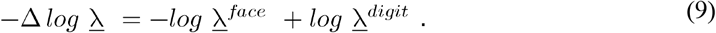

Due to the necessity of considering the interaction between signal rotation and manifold warping, their contribution must be evaluated together:

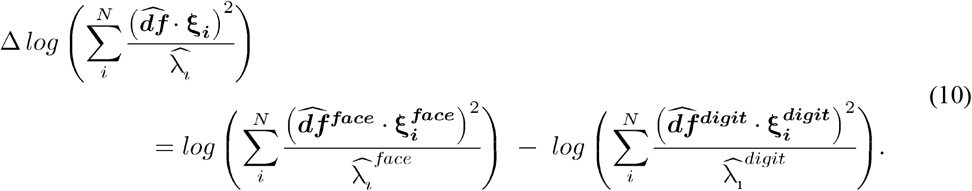

For each participant and each ROI, each type of logarithmic LFI (i.e., *log LFI*^*digit*^, *log LFI*^*SE*^, *log LFI* ^*MS*^, *log LFI*^*SR,MW*^, and *log LFI*^*face*^ was computed using responses to any position pair under two tasks and then averaged across position pairs within each distance bin. For each LFI type, each distance condition, and each ROI, the averaged LFI was pooled across the corresponding distance bins and participants.

To further investigate the joint contributions of signal rotation and manifold warping, we split the last addable item on each PC. Specifically, we computed (1) the numerator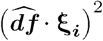 : the squared projected signal on each PC, (2) the denominator 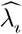: the relative variance on each PC, (3) the whole value 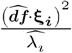 : the signal-to-noise ratio on each PC, and (4) the cumulative value 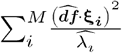 cumulative signal-to-noise ratio of the first *M* PCs. Each measure was averaged across position pairs within each distance bin. For each task, each distance condition, and each ROI, the mean was then pooled across the corresponding distance bins and participants.

### Statistical analysis

The statistical significance of the difference between two tasks or mechanisms was estimated using two-tailed sign tests when pooling voxels across all participants (i.e., for vRF properties, response variance and Fano factor) and Wilcoxon signed-rank tests for other cases. Bonferroni corrections were applied to account for multiple comparisons and we reported adjusted *p* values.

### Behavioral analysis

We calculated the d-prime^68^ to quantify participants’ sensitivity to the repetitions of face or digit identities in the main experiment. We set a response time window of 2 s after each repetition event. To ensure finite d-prime values, the minimum value of the number of misses or false alarms was set to 1. In the present study, the mean d-prime across participants was 3.032 ± 0.106 *S*.*D*. in the face task, and 3.650 ± 0.268 *S*.*D*. in the digit task. The behavioral results were shown in Supplementary Figure S8.

### Eye-tracking analysis

To exclude the possible confounding of eye movements in neural representations, we examined participants’ gazes in the main experiment. Preprocessing procedures for the eye-tracking data were followed those of Allen, et al. ^69^. Blinks were detected and removed within 100 ms before and 150 ms after each occurrence. Outliers that were more than 6° from the center of the screen were removed within 250 ms before and 250 ms after each occurrence. A linear drift trend was fitted and removed from the time series, and the gaze positions were then median-centered for each run. The time series data were downsampled to 100 Hz and smoothed with a 50 ms running average.

For each participant and each task, the gaze positions were segmented according to the time windows of valid trials and aggregated into 16 groups according to the face positions. We plotted the gaze positions for each face position group as a 2D histogram, then fitted a 2D Gaussian probability distribution and plotted the estimated 95% contours (see Supplementary Figure S9). The angles of the true face positions were calculated with respect to the x-axis, as were the angles of the major axes of the estimated gaze ellipses. Spearman’s correlation tests were performed on the 16 face position angles and the 16 gaze distribution angles for each participant and each task. The results showed that the gaze distributions cannot predict face position angles (*ps* > 0.05) across all participants in both tasks. This result suggests that participants were able to maintain central fixation in both the face and digit tasks without exhibiting a bias toward the presented face positions.

## ACKNOWLEDGMENT

This work was supported by the National Natural Science Foundation of China (32441102 to R.-Y. Z., 32371154 to Y.L.) and Shanghai Municipal Education Commission (2024AIZD014) to R.-Y.Z. Shanghai Rising-Star Program (24QA2705500) to Y.L..

## AUTHOR CONTRIBUTIONS

Y-Q. Y. and R-Y.Z. conceived and designed the study. Y-Q. Y. and S.L. conducted the experiment and preprocessed the fMRI data. Y.L. aided the data collection. K.K. prepared computer codes for several fMRI experiments and analyses. Y-Q. Y., K.K. and R-Y.Z. conducted the in-depth analysis of the fMRI data. Y-A., C. provided additional feedback on data analysis. Y-Q. Y. and R-Y.Z. wrote the first draft of the manuscript. All authors revised the manuscript.

## DATA AVAILABILITY

The data to reproduce the figure in this manuscript is available upon reasonable request.

## CODE AVAILABIBILTY

The code to reproduce the results in this manuscript is publicly available via GitHub at https://github.com/yqyou/facePRFattention.

## Supplementary Information for

## Table of Content

**Supplementary Figure S1**: The relationship between signal correlation and noise correlation.

**Supplementary Figure S2**: Supplementary results for four possible attention mechanisms in the near-distance condition.

**Supplementary Figure S3**: The effects of attentional mechanisms for the near-distance condition.

**Supplementary Figure S4**: The interaction effects between signal rotation and manifold warping.

**Supplementary Figure S5**: Eigen decomposition analyses for other ROIs in the far-distance condition.

**Supplementary Figure S6**: Eigen decomposition analyses for all ROIs in the near-distance condition.

**Supplementary Figure S7**: Effect of original and shuffled correlations on SVM decoding in the face task.

**Supplementary Figure S8**. The behavioral performance in the digit task and face task

**Supplementary Figure S9**: Eye tracking results exclude the confounding of eye movement

**Supplementary Note 1**: Derivation of the LFI formula based on eigen decomposition.

### Supplementary Figure S1

**Figure S1.**
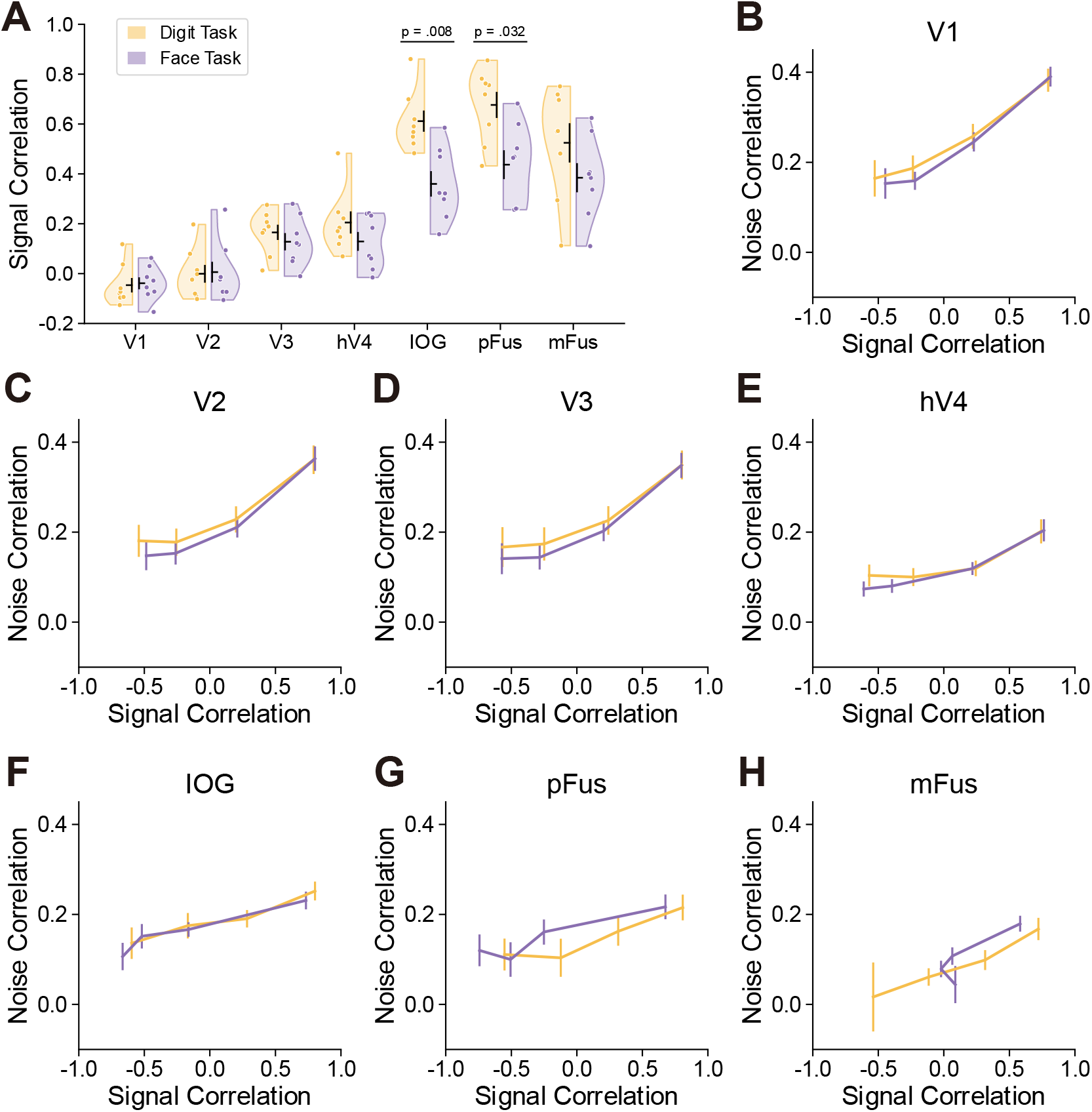
Relationship between signal correlation and noise correlation. ***A***. Signal correlation under two tasks for each ROI. Signal correlation was computed as the correlation between the averaged responses to 16 face positions for each pair of voxels. The median correlation across voxel pairs was taken. Each data point represents one participant. Two-tailed paired *t* tests on z-transformed signal correlation coefficients revealed significantly lower signal correlation in the face task for IOG (*p* = 0.008) and pFus (*p* = 0.032) after Bonferroni correction. ***B-H***. Noise correlation as a function of signal correlation under two tasks for each ROI. Voxel pairs were binned based on their signal correlation in the digit task (bin width = 0.5). Within each bin, the median signal correlation and noise correlation across voxel pairs were computed separately for each task. The error bars represent the mean ± S.E.M. across participants, and the lines are connected in bin order. A linear mixed model was applied to noise correlation, with task and bin as fixed factors and participants as random factor. The results showed a significant fixed effect of signal correlation bin in all ROIs (*ps* < 0.001), suggesting that the noise correlation significantly increased with signal correlation. The fixed effect of task was significant only in hV4 (*p* = 0.041), but not in other ROIs (*ps* > 0.116), suggesting that the noise correlation was significantly lower in face task compared to the digit task in hV4.

### Supplementary Figure S2

**Figure S2.**
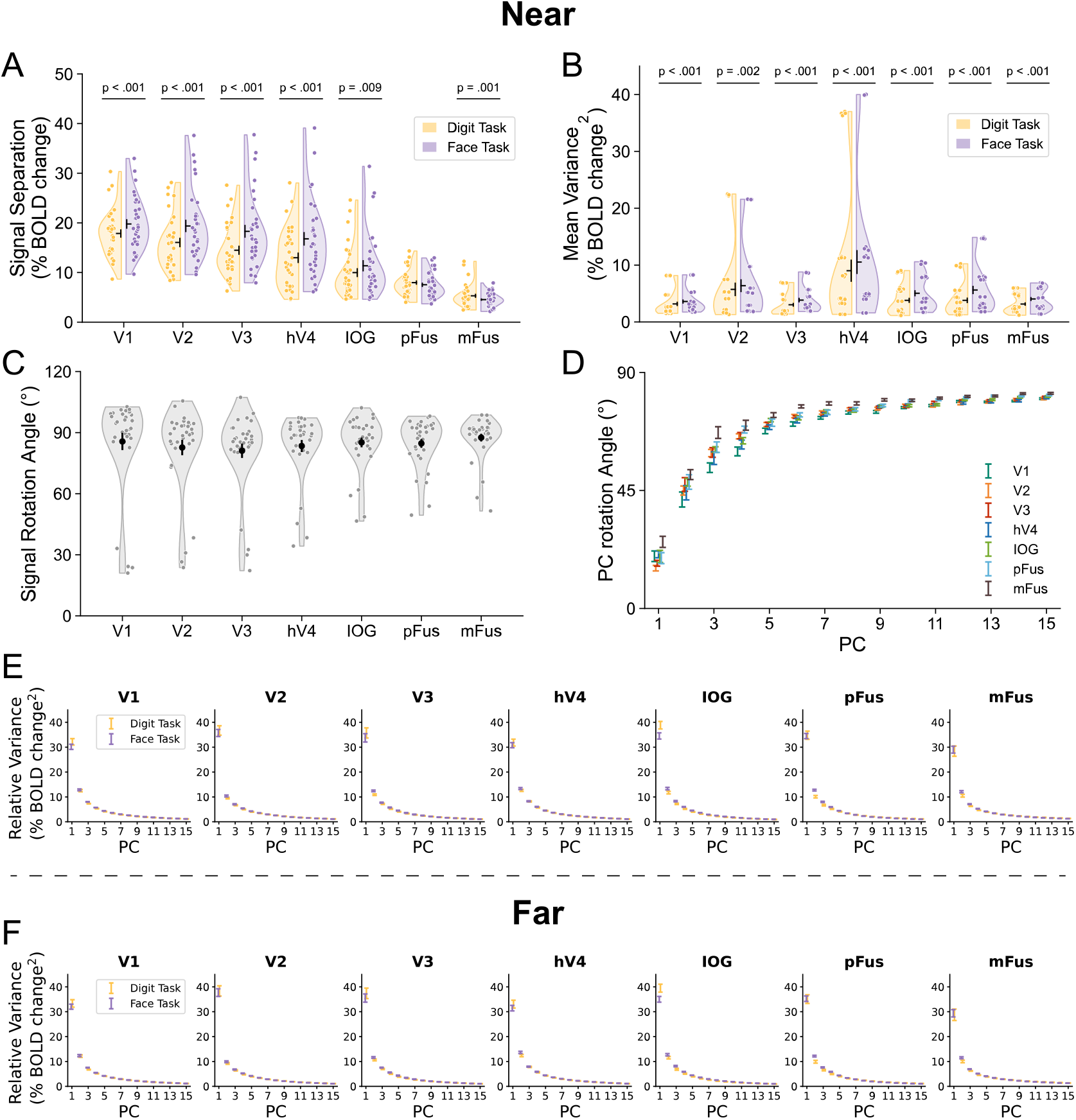
Supplementary results of four possible attention mechanisms at the population level. ***A****-****E***. The effects of attention on four possible mechanisms for the near distance condition in each ROI. ***F***. The effects of attention on eigenvalues of the mean covariance for the far distance condition in each ROI. Each data point represents a sample (8 participants × 4 distance bins = 32 samples) and the error bars represent mean ± S.E.M. across samples. All significant differences are labelled, with three asterisks indicating *p* < 0.001 (***A&B***).

### Supplementary Figure S3

**Figure S3.**
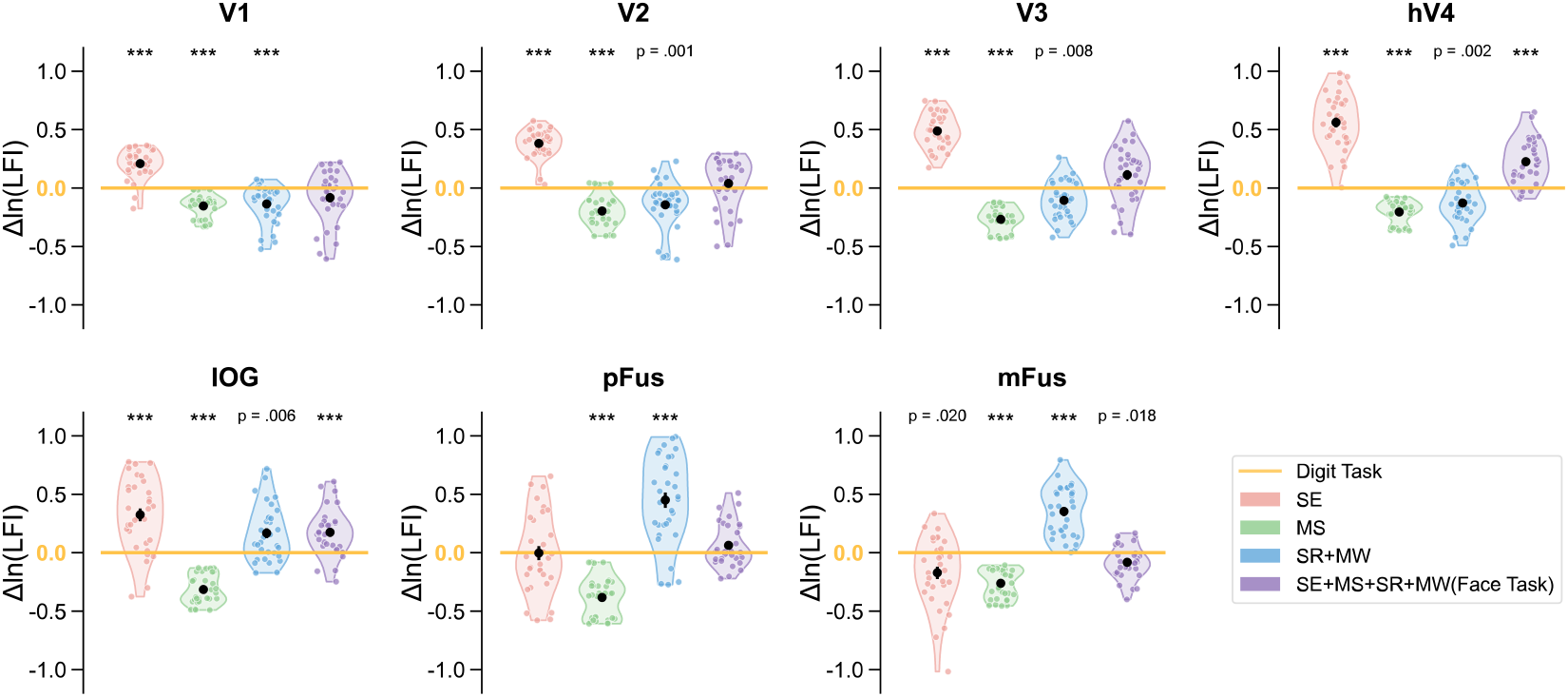
The difference between the LFI applying difference mechanisms and the LFI of the digit task in each ROI for the near-distance condition. Each data point represents a sample (8 participants × 4 distance bins = 32 samples) and the error bars represent mean ± S.E.M. across samples. All significant differences are labelled, with three asterisks indicating *p* < 0.001.

### Supplementary Figure S4

**Figure S4.**
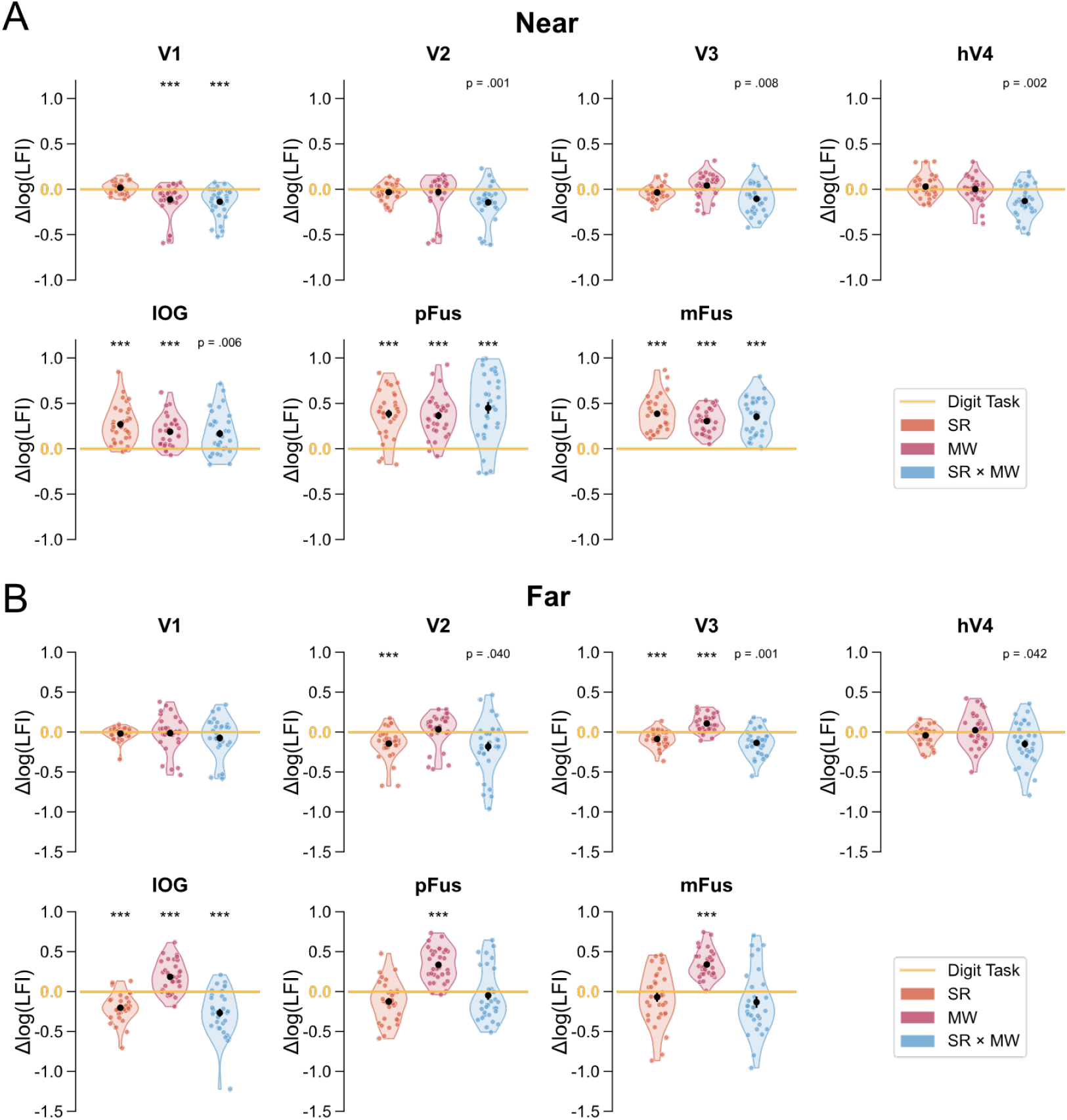
Differences in LFI relative to the digit task, resulting from applying signal rotation only, manifold warping only, or both. The results reveal a complex interaction between SR and MW rather than a simple additive effect. Data are shown for each ROI under the near-distance condition in panel (A) and the far-distance condition in panel (B). Each data point represents a sample (8 participants × 4 distance bins = 32 samples) and the error bars represent mean ± S.E.M. across samples. All significant differences are labelled, with three asterisks indicating *p* < 0.001.

### Supplementary Figure S5

**Figure S5.**
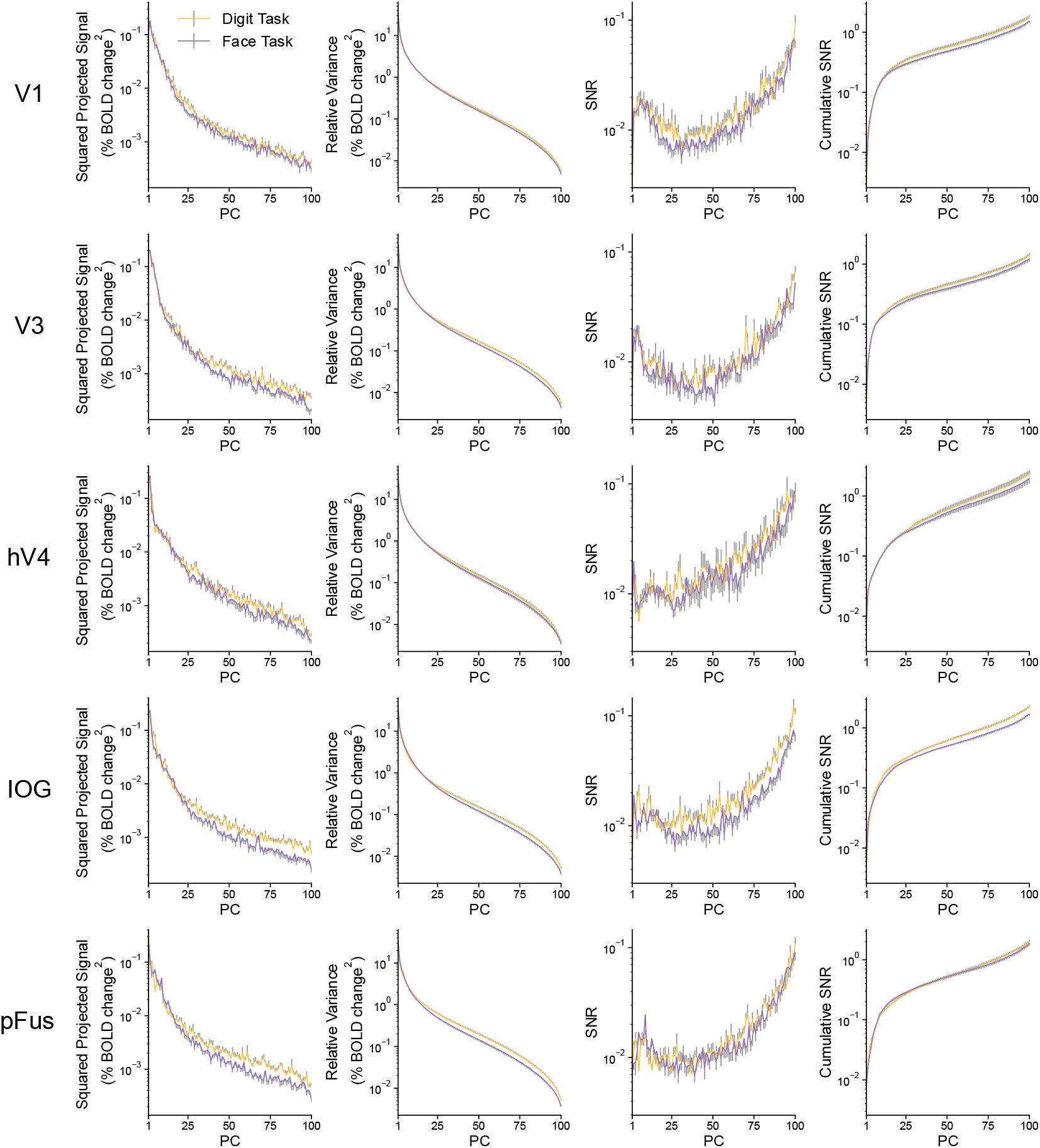
Eigen-decomposition analyses for the joint effects of signal rotation and manifold warping in V1, V3, hV4, IOG, and pFus for the far-distance condition. The x-axes are PCs ranked by eigenvalues from high to low. Each figure from left to right represented the squared projected signal 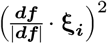 on each PC, the relative variance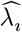 on each PC, the signal-to-noise ratio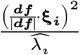 on each PC, and cumulative signal-to-noise ratio 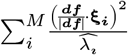 of the first *M* PCs. The mean ± S.E.M. across samples (8 participants × 4 distance bins = 32 samples) were represented by error bars, where only half of the error bars were drawn for better presentation.

### Supplementary Figure S6

**Figure S6.**
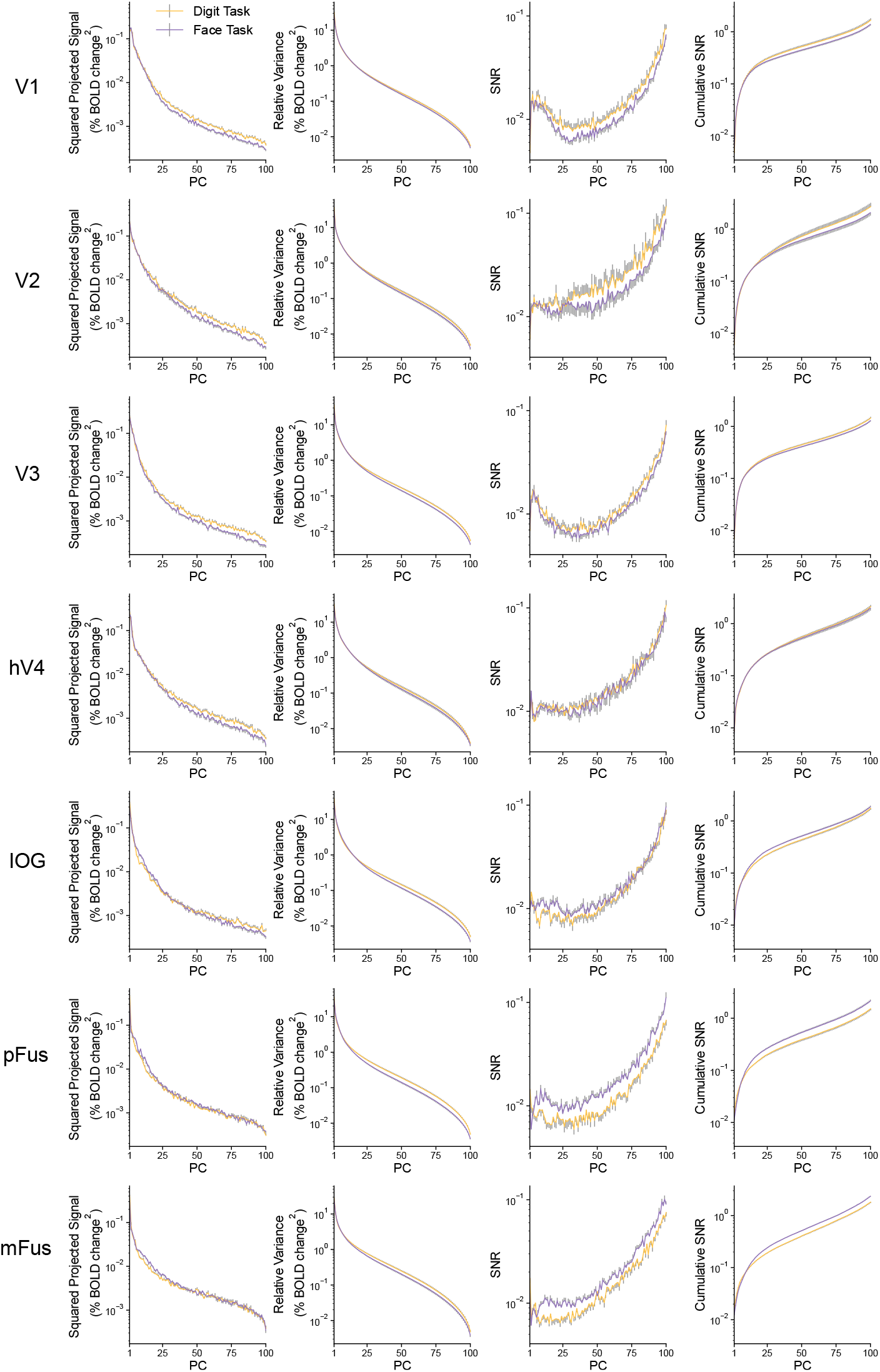
Eigen-decomposition analyses for the joint effects of signal rotation and manifold warping in all ROIs for the near-distance condition. The x-axes are PCs ranked by eigenvalues from high to low. Each figure from left to right represented the squared projected signal (***df*** · *ξ* _*i*_)^2^on each PC, the relative variance 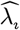 on each PC, the signal-to-noise ratio 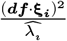 on each PC, and cumulative signal-to-noise ratio 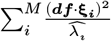 of the first *M* PCs. Each data point indicates a sample (8 participants × 16 positions = 128 samples). The mean ± S.E.M. were represented by error bars, where only half of the error bars were drawn for better presentation.

### Supplementary Figure S7

**Figure S7:**
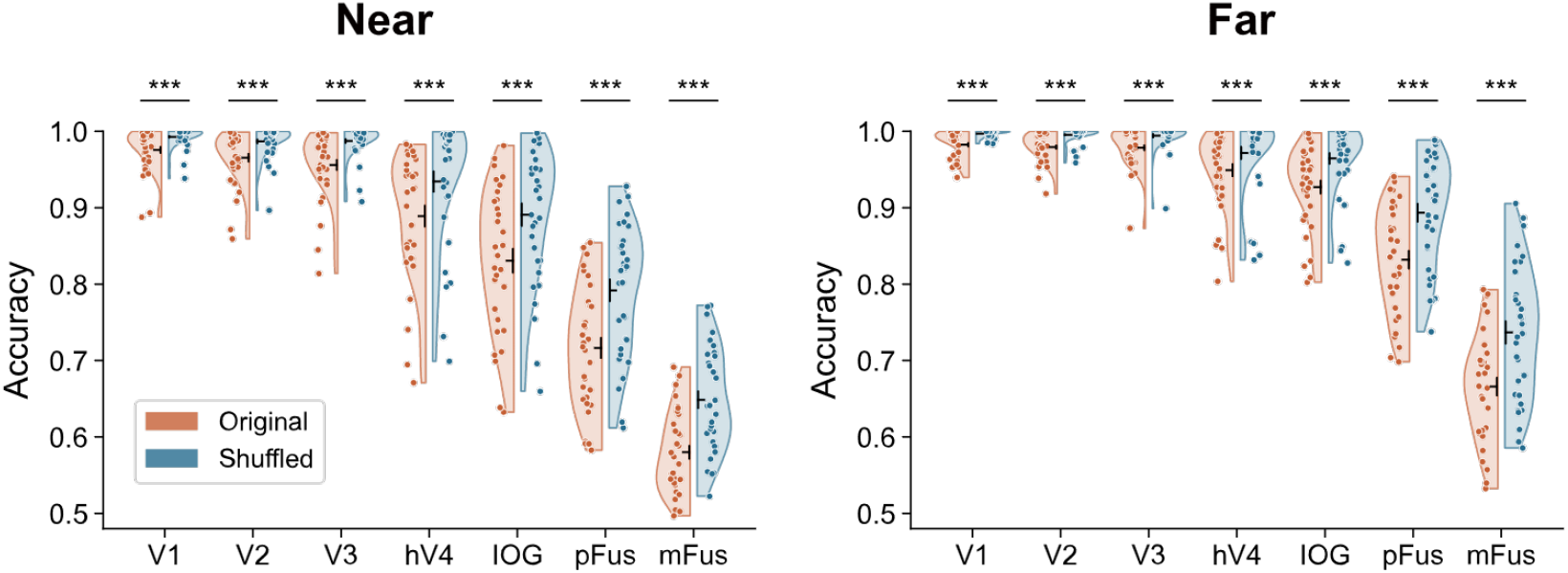
Effect of original and shuffled correlations on SVM decoding in the face task. Linear SVM decoding accuracy is shown for shuffled and original voxel responses in each task and ROI. Following a procedure similar to Jiang, et al. ^1^, the correlations were disrupted by shuffling the trial order independently for each voxel. This procedure was repeated 1,000 times to generate a distribution. The distribution for the original dataset was obtained by shuffling trial order consistently across voxels for each repeat. Each data point indicates a sample (8 participants × 4 distance bins = 32 samples). The black horizontal and vertical lines in the violin plots represent the mean and ± S.E.M. across samples, respectively. All significant differences are labeled, with three asterisks indicating *p* < 0.001.

### Supplementary Figure S8

**Figure S8.**
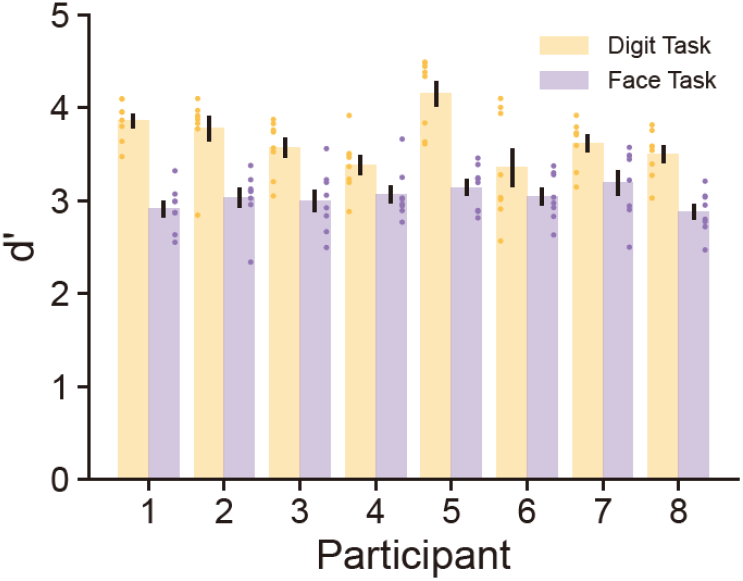
The mean d-prime during the digit task and face task for each participant. Each data point represents the d-prime for one session of one participant and the error bars represent mean ± S.E.M. across 8 sessions.

### Supplementary Figure S9

**Figure S9.**
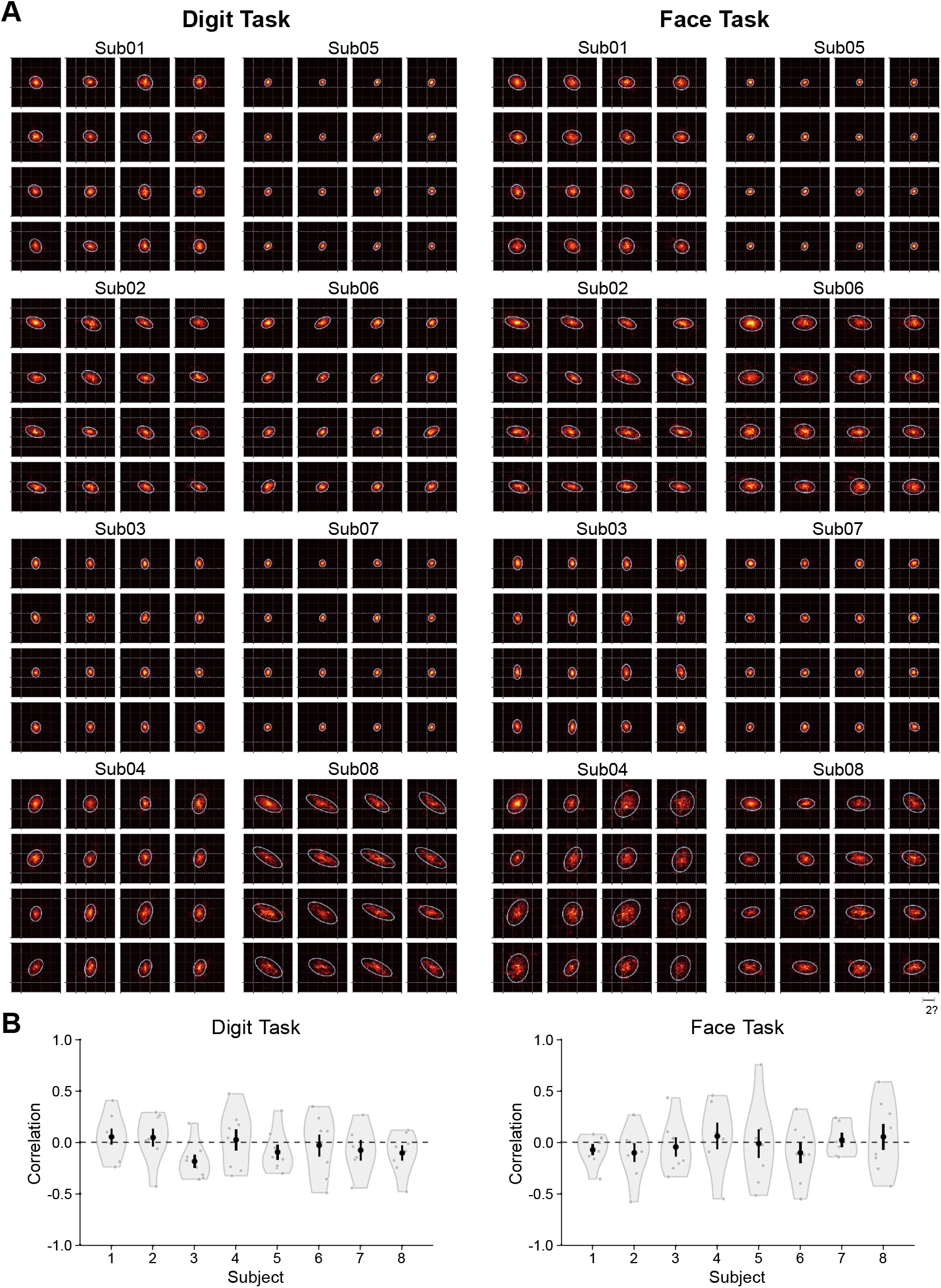
Eye tracking results for each participant during the digit and face tasks. ***A***. Each image is a 2D histogram of gaze positions recorded during the presentation of faces at a single location within a 4 × 4 grid. Horizontal and vertical gray lines represent 2° increments, with the total range spanning from -5° to 5°. The white contour encloses 95% of the fitted 2D Gaussian distribution of gaze positions. The concentration of gaze distributions varied across participants due to measurement noise from the instrument. However, regardless of the face position, the gaze distribution consistently remained centered on the screen, and the shape of the distribution did not exhibit any bias toward the face location. ***B***. Correlation between the angle of the principal axis of the gaze distribution and the angle of the face position relative to the horizontal direction, across 16 face positions. Each data point represents the correlation for one session of one participant. Two-tailed one-sample t-tests were performed, and the results showed that the correlations for all participants were not significantly different from zero in both tasks after Bonferroni correction. Each data point represents one participant and the error bars represent mean ± S.E.M. across participants.

## Supplementary Note 1: Derivation of the LFI formula based on eigen decomposition

The original formular of the LFI was given as follows:

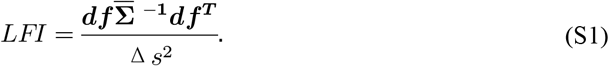

We performed eigen decomposition to the mean covariance 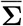:

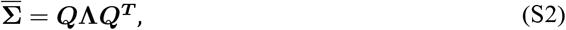

where ***Q*** denotes the orthogonal matrix (ℝ^*N*×*N*^) whose columns are mutually orthogonal eigenvectors [***ξ***_**1**_, ***ξ***_**2**_, …, ***ξ***_***i***_], and **Λ** denotes the diagonal matrix (ℝ^*N*×*N*^) containing the eigenvalues *diag* (*λ*_1_, *λ*_2_, …, *λ*_*i*_) corresponding to the eigenvectors.

The LFI formula can be rewritten as:

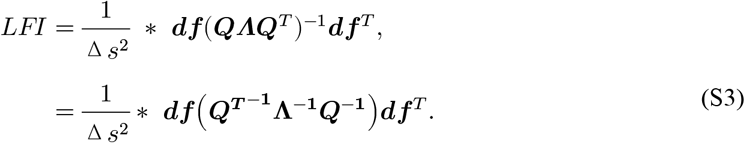

Since ***Q*** is the orthogonal matrix, where ***Q***^***T***^ = ***Q***^−_**1**_^, so

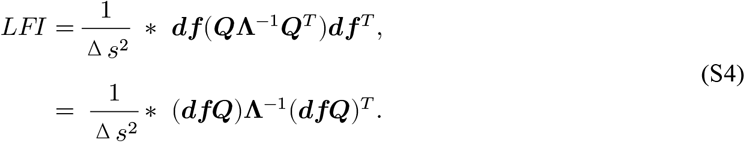

(***dfQ***) gives a 1 × *N* vector, where the *i*-th value is ***df*** ⋅ ***ξ***_*i*_.

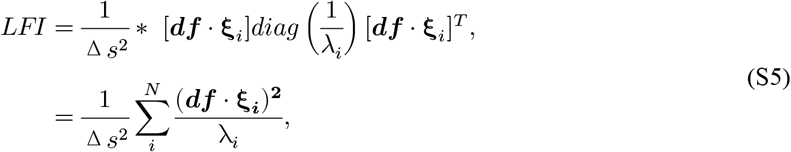

We represented the signal vector ***df*** with its modulus length |***df***| and the unit vector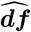, and split *i*-th eigenvalue *λ*_i_ into the mean of the eigenvalues 𝜆 and the relative eigenvalue 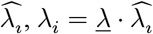. The final formula was expressed as:

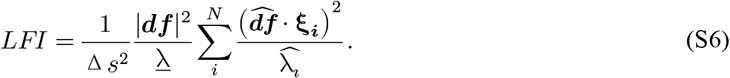

We take the log on both sides of Eq. S6:

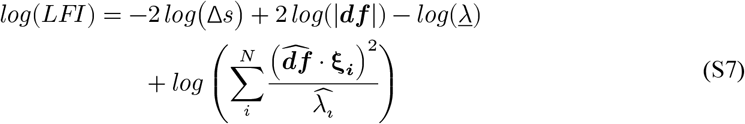

